# Pre-existing and *de novo* humoral immunity to SARS-CoV-2 in humans

**DOI:** 10.1101/2020.05.14.095414

**Authors:** Kevin W. Ng, Nikhil Faulkner, Georgina H. Cornish, Annachiara Rosa, Ruth Harvey, Saira Hussain, Rachel Ulferts, Christopher Earl, Antoni Wrobel, Donald Benton, Chloe Roustan, William Bolland, Rachael Thompson, Ana Agua-Doce, Philip Hobson, Judith Heaney, Hannah Rickman, Stavroula Paraskevopoulou, Catherine F. Houlihan, Kirsty Thomson, Emilie Sanchez, David Brealey, Gee Yen Shin, Moira J. Spyer, Dhira Joshi, Nicola O’Reilly, Philip A. Walker, Svend Kjaer, Andrew Riddell, Catherine Moore, Bethany R. Jebson, Meredyth G.Ll. Wilkinson, Lucy R. Marshall, Elizabeth C. Rosser, Anna Radziszewska, Hannah Peckham, Coziana Ciurtin, Lucy R. Wedderburn, Rupert Beale, Charles Swanton, Sonia Gandhi, Brigitta Stockinger, John McCauley, Steve Gamblin, Laura E. McCoy, Peter Cherepanov, Eleni Nastouli, George Kassiotis

## Abstract

Several related human coronaviruses (HCoVs) are endemic in the human population, causing mild respiratory infections^1^. Severe Acute Respiratory Syndrome Coronavirus 2 (SARS-CoV-2), the etiologic agent of Coronavirus disease 2019 (COVID-19), is a recent zoonotic infection that has quickly reached pandemic proportions^2,3^. Zoonotic introduction of novel coronaviruses is thought to occur in the absence of pre-existing immunity in the target human population. Using diverse assays for detection of antibodies reactive with the SARS-CoV-2 spike (S) glycoprotein, we demonstrate the presence of pre-existing humoral immunity in uninfected and unexposed humans to the new coronavirus. SARS-CoV-2 S-reactive antibodies were readily detectable by a sensitive flow cytometry-based method in SARS-CoV-2-uninfected individuals and were particularly prevalent in children and adolescents. These were predominantly of the IgG class and targeted the S2 subunit. In contrast, SARS-CoV-2 infection induced higher titres of SARS-CoV-2 S-reactive IgG antibodies, targeting both the S1 and S2 subunits, as well as concomitant IgM and IgA antibodies, lasting throughout the observation period of 6 weeks since symptoms onset. SARS-CoV-2-uninfected donor sera also variably reacted with SARS-CoV-2 S and nucleoprotein (N), but not with the S1 subunit or the receptor binding domain (RBD) of S on standard enzyme immunoassays. Notably, SARS-CoV-2-uninfected donor sera exhibited specific neutralising activity against SARS-CoV-2 and SARS-CoV-2 S pseudotypes, according to levels of SARS-CoV-2 S-binding IgG and with efficiencies comparable to those of COVID-19 patient sera. Distinguishing pre-existing and *de novo* antibody responses to SARS-CoV-2 will be critical for our understanding of susceptibility to and the natural course of SARS-CoV-2 infection.

## Results

Immune cross-reactivity among seasonally spreading human coronaviruses (HCoVs) has long been hypothesised to provide cross-protection, albeit transient, against infection with distinct HCoV types^1,4,5^. To determine the degree of cross-reactivity between HCoVs and the recently introduced zoonotic coronavirus SARS-CoV-2, we developed a sensitive flow cytometry-based assay for detection of SARS-CoV-2-binding antibodies (Methods). The main target for such antibodies is the spike (S) glycoprotein, which is proteolytically processed into the S1 and S2 subunits, mediating target cell attachment and entry, respectively^6,7^. To ensure detection of antibodies targeting all available epitopes in naturally processed S, we transfected HEK293T cells with an expression vector encoding the wild-type S of SARS-CoV-2 and stained them with primary serum samples.

Staining with the S1-reactive CR3022 monoclonal antibody (IgG class) identified a smaller percentage of SARS-CoV-2 S-expressing HEK293T cells and was of lower intensity than staining with COVID-19 convalescent sera (Extended data Fig. 1), indicating that polyclonal IgG antibodies targeted a wider range of epitopes presented on these cells. As well as IgG antibodies, this assay readily detected SARS-CoV-2 S-reactive IgM and IgA antibodies in COVID-19 convalescent sera (Extended data Fig. 2). Consistent with prior reports^8-18^, titres of SARS-CoV-2 S-reactive antibodies varied by several orders of magnitude between COVID-19 convalescent sera, with some reliably detected only at the highest serum concentration (1:50 dilution) (Extended data Fig. 2), which was used for subsequent sensitive measurements. Whilst the intensity of IgM and IgG staining decreased proportionally with serum dilutions, the intensity of IgA staining peaked at intermediate serum dilutions (Extended data Fig. 2), likely owing to competition with IgG antibodies at higher serum concentrations. Therefore, the intensity of staining with each of the three main Ig classes might reflect their relative ratios, as well as their absolute titres. In contrast, the percentage of cells stained with each Ig class was less sensitive to serum dilutions and was chosen as a correlate for seropositivity. Indeed, the percentage of SARS-CoV-2 S-transfected cells stained with all three Ig classes (IgM^+^IgG^+^IgA^+^) distinguished COVID-19 sera from control sera with a high degree of sensitivity and specificity (Extended data Fig. 3a-c). A total of 170 RT-qPCR-confirmed COVID-19 patients at University College London Hospitals (UCLH) and 262 SARS-CoV-2-uninfected patients and healthy donors (Table S1) were tested by the flow cytometry-based assay, demonstrating overall 91.7% sensitivity and 100% specificity of seropositivity detection (Extended data Fig. 3b). All but one of the antibody-negative COVID-19 patient samples were collected between days 2 and 16 post symptoms onset (Extended data Fig. 3d). Consequently, for samples taken after day 16 post symptoms onset, the flow cytometry-based assay demonstrated 99.8% sensitivity and 100% specificity of seropositivity detection, with no evidence of decline at the latest time-point examined (day 43 post onset of symptoms) (Extended data Fig. 3c).

The single COVID-19 patient without a concurrent IgM, IgG and IgA response on day 29 post symptoms onset, when all other COVID-19 patients had seroconverted, did exhibit SARS-CoV-2 S-reactive IgG antibodies detected by this assay in the absence of the other two Ig classes (Extended data Fig. 3d). Inspection of patient records indicated that this was a 60-year-old bone marrow transplant recipient, who tested positive for HCoV infection in February 2020 and subsequently contracted SARS-CoV-2, testing positive by RT-qPCR the following month (Extended data Fig. 4).

Serum samples taken around the diagnosis of HCoV and SARS-CoV-2 infections were negative for SARS-CoV-2 S-reactive antibodies, likely due to the immunosuppressed state of this patient (Extended data Fig. 4). However, serum taken 3 weeks later contained SARS-CoV-2 S-reactive IgG, in the absence of other antibody classes (Extended data Fig. 4). This profile was more consistent with an anamnestic response to HCoVs, than a *de novo* response to SARS-CoV-2. This patient described only mild COVID-19 symptoms that did not necessitate hospitalisation, but appeared chronically infected with SARS-CoV-2, having tested repeatedly positive for viral RNA for over a month (Extended data Fig. 4).

To examine whether SARS-CoV-2 S-reactive IgG antibodies, detectable by the FACS-based assay, might have been induced by prior HCoV infection, we tested sera from SARS-CoV-2-uninfected patients with or without RT-qPCR-confirmed HCoV infection, sampled before or during the early spread of SARS-CoV-2 in the UK (Table S1). Although the presence of SARS-CoV-2 S-reactive antibodies of all three Ig classes effectively distinguished COVID-19 patients, remarkably, a proportion of SARS-CoV-2-uninfected patients also had SARS-CoV-2 S-specific antibodies (Fig. 1a). Again, sera from the latter group contained predominantly lower levels of S-reactive IgG and no IgM or IgA antibodies (Fig. 1a), suggesting the presence of cross-reactive immunological memory.

**Figure 1.**
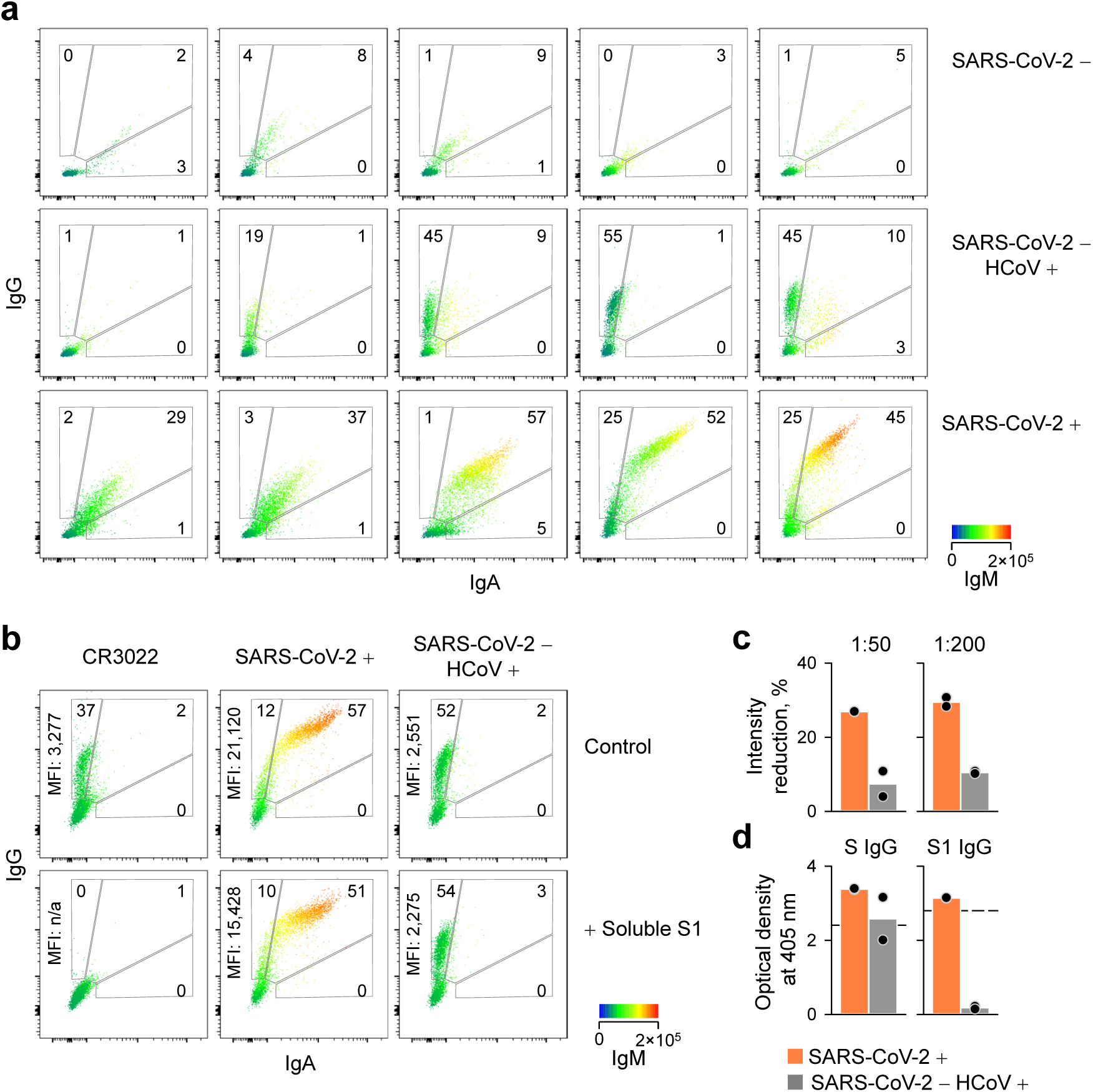
Flow cytometric detection and specificity of antibodies reactive with SARS-CoV-2 S. **a**, FACS profiles of IgG, IgA and IgM antibodies in 5 individual patient sera from each indicated group. Levels of IgM are indicated by a heatmap. **b**, Ability of soluble S1 to inhibit binding to SARS-CoV-2 S of the S1-specific CR3022 antibody or antibodies in the sera of SARS-CoV-2-infected (SARS-CoV-2 +) or HCoV-infected (SARS-CoV-2 - HCoV +) patients. One representative of two patients is shown. **c**, Quantitation of the inhibitory effect of soluble S1 on binding to SARS-CoV-2 S of sera from patients described in b. Each dot is an individual patient serum tested at 1:50 and 1:200 dilution. **d**, Optical densities from ELISAs coated with S or S1 of sera from patients described in b. Each dot is an individual patient serum tested at 1:50 dilution. The dashed lines represents the optical densities obtained with the CR3022 antibody.

The S2 subunit exhibits a higher degree of homology among coronaviruses than S1 (Extended data Fig. 5) and was likely the main target of cross-reactive antibodies^6^. Indeed, whereas the addition of recombinant soluble S1 completely abolished binding of the S1-reactive CR3022 antibody^19^ to SARS-CoV-2 S-expressing cells, it did not affect the binding of HCoV patient sera (Fig. 1b,c). Binding of COVID-19 patient sera was reduced by approximately 30% by soluble S1, indicating recognition of both S1 and S2 (Fig. 1b,c). Consistent with these observations, sera from both COVID-19 and HCoV patients reacted comparably in a SARS-CoV-2 S-coated ELISA, whereas only those from COVID-19 patients reacted in a SARS-CoV-2 S1-coated ELISA (Fig. 1d). These data highlighted cross-reactivity of HCoV patient sera with SARS-CoV-2 S, and specific recognition of the S1 subunit by COVID-19 patient sera.

To confirm and determine the extent of such cross-reactivity using an independent assay, we developed ELISAs with a number of SARS-CoV-2 recombinant proteins (Methods). These included the stabilised trimeric S ectodomain, the S1 subunit, the receptor-binding domain (RBD) and the nucleoprotein (N). In a cohort of 135 COVID-19 patients (Table S1), the sensitivity of ELISAs was lower than that of the FACS-based assay, with 20 COVID-19 samples testing negative on ELISA coated with the S1 subunit, 8 of which also tested negative on S-coated or N-coated ELISAs (Extended data Fig. 6). Overall, the S1-coated ELISA for IgG detection demonstrated 92.1% sensitivity and 99.6% specificity of seropositivity detection for all samples taken after day 16 post symptoms onset (Extended data Fig. 7a-c). Rates of IgG seropositivity determined by SARS-CoV-2 S1-coated ELISA were congruent with, but overall lower than those determined by FACS, underscoring the higher sensitivity of the FACS-based assay (Extended data Fig. 7d,e).

We next tested our initial cohort of COVID-19 patients and SARS-CoV-2-uninfected individuals (SARS-CoV-2 -), which included earlier samples of unknown HCoV status and more recent samples of known HCoV status (Table S1), in the different assays. SARS-CoV-2 S-reactive IgG antibodies were detected by FACS in 5 of 34 SARS-CoV-2-uninfected individuals with recent HCoV infection, as well as in 1 of 31 individuals without recent HCoV infection (Fig. 2a). This suggested that these antibodies might have persisted from earlier HCoV infections, rather than having been induced by the most recent one. The 3 SARS-CoV-2-uninfected individuals with the highest cross-recognition of S on FACS, and a further 4 individuals, also had IgG antibodies detectable on ELISA coated with SARS-CoV-2 S ectodomain, as well as with SARS-CoV-2 N (Fig. 2b-e), also highly conserved among coronaviruses (CoVs). In contrast, none of the control samples had IgG antibodies detectable on ELISAs coated with the less conserved SARS-CoV-2 S1 or RBD (Fig. 2c,e). Collectively, these results suggested that a total of 12 of 95 control samples exhibited IgG antibodies cross-reactive with conserved epitopes in SARS-CoV-2 proteins (S2 and N). Moreover, a sample from an 81-year-old COVID-19 patient in this cohort collected on day 16 post mild COVID-19 symptoms exhibited only IgG reactivity to SARS-CoV-2 S, but not to S1, which would be more characteristic of pre-existing antibody memory to HCoVs, than a *de novo* response to SARS-CoV-2 (Fig. 2a-e).

**Figure 2.**
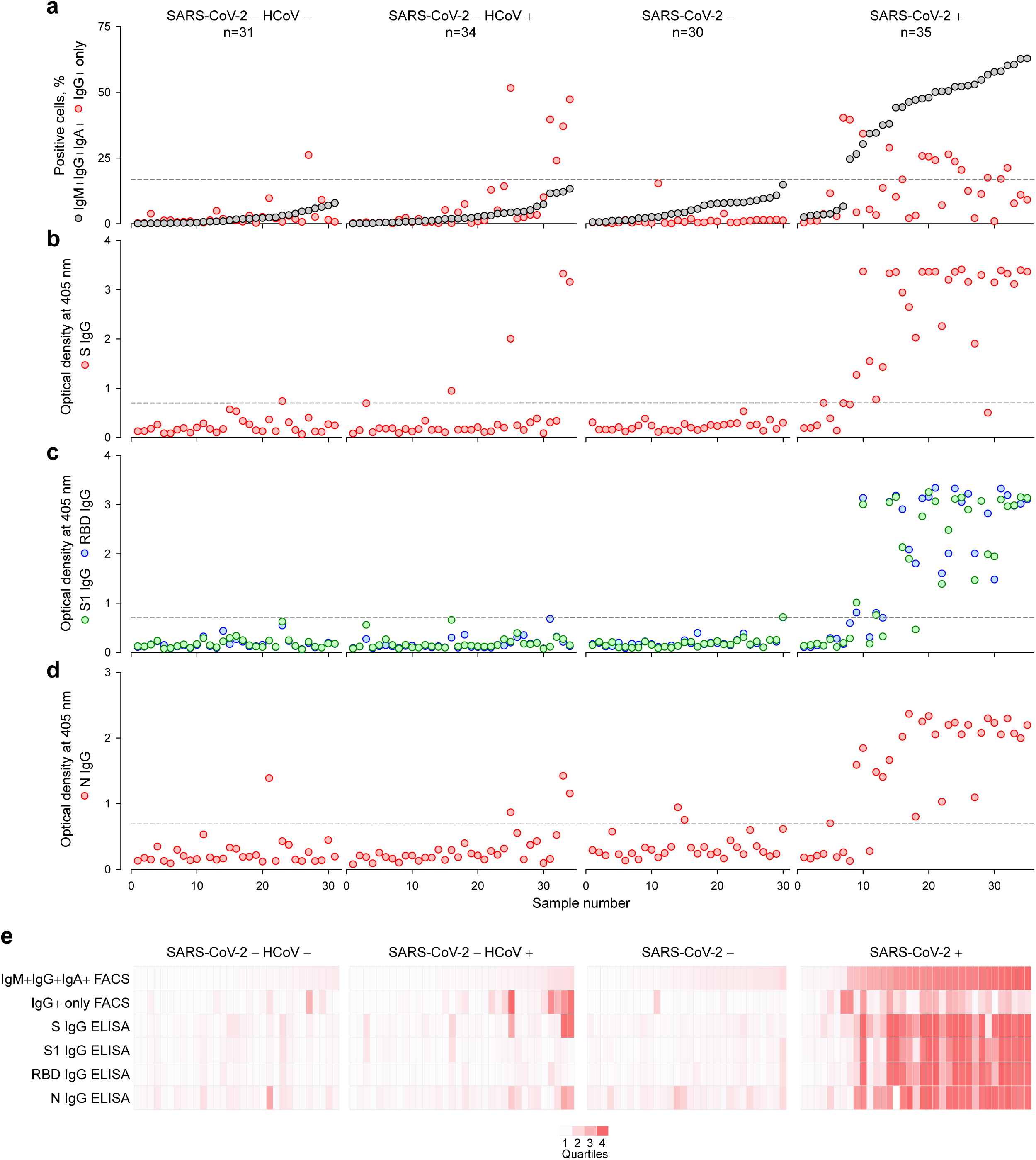
Comparison of antibody detection methods using a panel of patient samples. Serum samples from the following groups were compared in all panels: SARS-CoV-2-uninfected without recent HCoV infection (SARS-CoV-2 - HCoV -); SARS-CoV-2-uninfected with recent HCoV infection (SARS-CoV-2 - HCoV +); SARS-CoV-2-uninfected with unknown history of recent HCoV infection (SARS-CoV-2 -); SARS-CoV-2-infected (SARS-CoV-2 +). **a**, Frequency of cells that stained with all three antibody classes (IgM+IgG+IgA+) or only with IgG (IgG+) in each of these samples, ranked by their IgM+IgG+IgA+ frequency. **b-d**, Optical densities from ELISAs coated with S (b), S1 or RBD (c) or N (d) of the same samples. Dashed lines in a-d denote the assay sensitivity cut-offs. **e**, Summary of the results from a-d, represented as a heatmap of the quartile values.

Although the data presented here and emerging in the recent and unrefereed literature^14,20,21^ suggest extensive antibody cross-reactivity between SARS-CoV-2 and HCoVs, the potential consequences of pre-existing immunity for the course of SARS-CoV-2 infection or susceptibility to COVID-19 have not been fully considered. One of the ways in which HCoV-elicited antibodies might protect against SARS-CoV-2 infection is inhibition of entry into the target cell. SARS-CoV-2 binding to one identified cellular receptor, angiotensin-converting enzyme 2 (ACE2)^2,6,7,22^, is mediated by the RBD. As HCoV-induced antibodies cross-react with S2, but not the RBD, they might not be expected to block ACE2-RBD interactions. However, SARS-CoV-2-neutralising antibodies that do not block RBD-ACE2 interaction have been reported^23,24^, as have neutralising antibodies targeting the S2 subunit of SARS-CoV^25,26^ or of SARS-CoV-2^24^. It is therefore plausible that HCoV patient sera targeting the S2 also neutralise, either by indirectly affecting ACE2-mediated viral entry or by interfering with alternative routes of entry. Indeed, entry of SARS-CoV-2 can also be facilitated by the alternative receptor CD147, also known as basigin (BSG)^27^, neuropilin 1^28,29^, and possibly also by receptor-independent mechanisms, as has been described for other CoVs^30,31^. To examine the ability of HCoV patient sera to inhibit viral entry, we used HEK293T cells as targets. These cells lack ACE2 expression, but are nevertheless permissive to entry of lentiviral particles pseudotyped with SARS-CoV-2 S (Extended data Fig. 8). Moreover, transduction efficiency of HEK293T cells by SARS-CoV-2 S pseudotypes was not further increased by ACE2 overexpression (Extended data Fig. 8), highlighting ACE2-independent entry. In contrast, HEK293T cells expressed high levels of *BSG* and *NRP1*, encoding CD147 and neuropilin 1, respectively (Extended data Fig. 8).

Sera from seroconverted (Ab +) COVID-19 patients efficiently neutralised SARS-CoV-2 S pseudotypes, consistent with prior reports^10,15,17,32,33^, whereas those from COVID-19 patients that had not yet produced binding antibodies (Ab -) did not (Fig. 3a). Surprisingly, sera from SARS-CoV-2-uninfected patients that contained SARS-CoV-2 S-reactive antibodies neutralised these pseudotypes, with efficiencies comparable with sera from seroconverted COVID-19 patients (Fig. 3a). None of the sera neutralised lentiviral particles pseudotyped with the control glycoprotein of vesicular stomatitis virus (VSV) (Fig. 3a). Titres of neutralising antibodies in COVID-19 patient sera correlated well with titres of SARS-CoV-2 RBD-binding antibodies, determined by ELISA and with titres of SARS-CoV-2 S-binding antibodies, determined by FACS (Fig. 3b). Despite the low numbers, the correlation between neutralising antibodies in SARS-CoV-2-uninfected patient sera with SARS-CoV-2 S-binding IgG antibodies determined by FACS was even stronger (Fig. 3c). Collectively, these data indicated that HCoV-elicited cross-reactive antibodies can interfere with at least one mode of SARS-CoV-2 entry into target cells. To confirm these findings in different experimental setting, we used infection of Vero E6 cells with authentic SARS-CoV-2 and assessed the ability of patient sera to reduce the number and size of plaques formed. All COVID-19 patient sera tested had measurable neutralising activity in this assay (Fig. 3d). Importantly, 5 of 6 sera from SARS-CoV-2-uninfected patients with FACS-detectable SARS-CoV-2 S-reactive antibodies also neutralised SARS-CoV-2 in this assay, albeit on average less potently than COVID-19 patient sera, whereas none of the sera from SARS-CoV-2-uninfected patients from the same cohort, but without FACS-detectable SARS-CoV-2 S-reactive antibodies did (Fig. 3d), supporting the functional relevance of FACS-detectable SARS-CoV-2 S-binding IgG antibodies.

**Figure 3.**
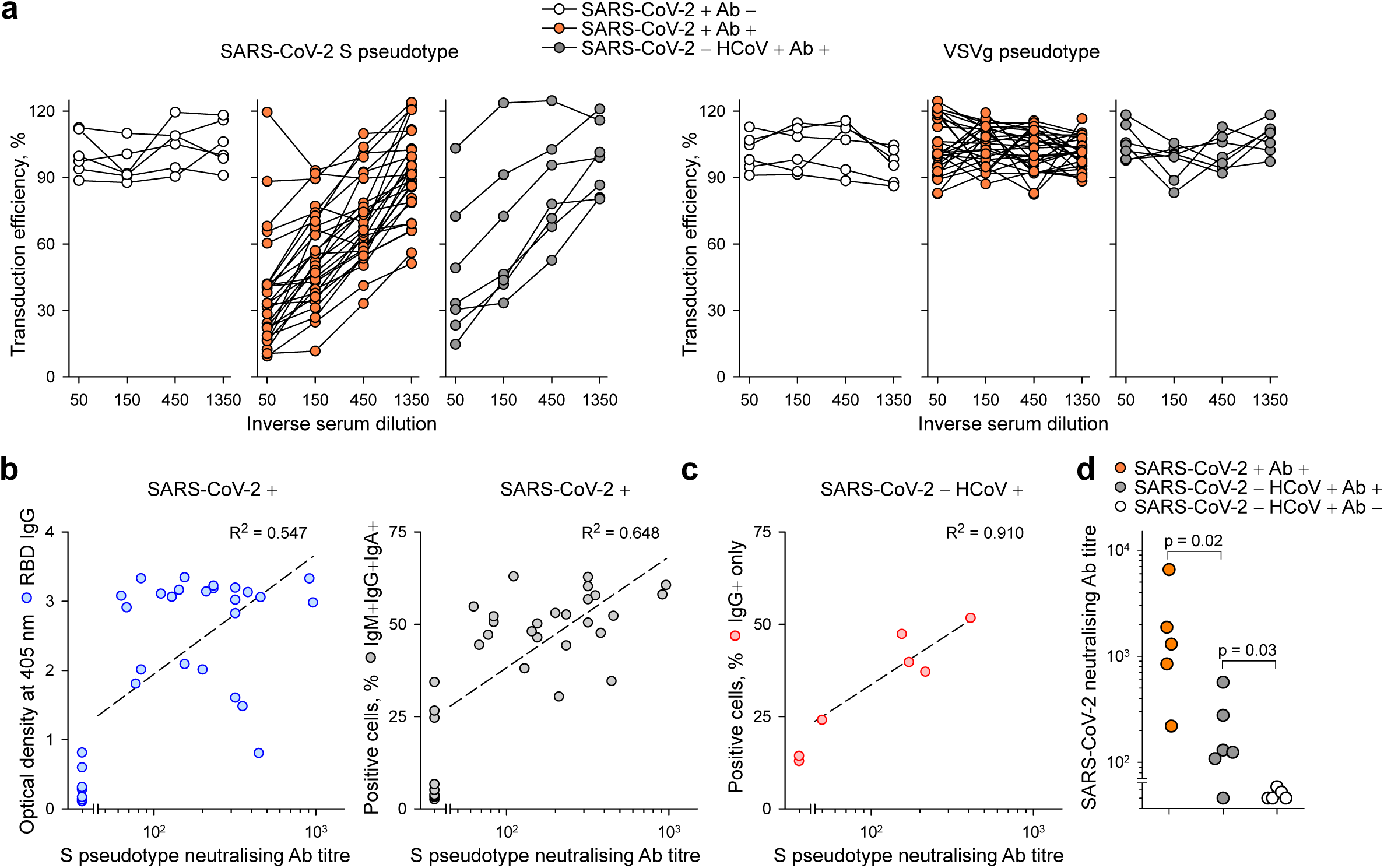
Neutralisation of SARS-CoV-2 S pseudotypes and authentic SARS-CoV-2 by SARS-CoV-2-infected and -uninfected patient sera. **a**, Percent inhibition of transduction efficiency of SARS-CoV-2 S (*left*) or control VSVg (*right*) lentiviral pseudotypes by SARS-CoV-2-infected patient sera without (SARS-CoV-2 + Ab -) or with (SARS-CoV-2 + Ab +) detectable SARS-CoV-2 S-binding antibodies and those from SARS-CoV-2-uninfected patients with recent HCoV infection and with detectable SARS-CoV-2 S-binding antibodies (SARS-CoV-2 - HCoV + Ab +). Each line is an individual serum sample. **b**, Correlation of neutralising antibody titres in SARS-CoV-2-infected patient sera with optical densities from RBD-coated ELISAs (*left*) or IgM+IgG+IgA+ staining frequencies, determined by FACS (*right*). **c**, Correlation of neutralising antibody titres in sera from SARS-CoV-2-uninfected patients with recent HCoV infection with IgG+ staining frequencies, determined by FACS. **d**, Authentic SARS-CoV-2 neutralisation titres of sera from seroconverted COVID-19 patients (SARS-CoV-2 + Ab +) or from SARS-CoV-2-uninfected patients with recent HCoV infection and with (SARS-CoV-2 - HCoV + Ab +) or without (SARS-CoV-2 - HCoV + Ab -). FACS-detectable SARS-CoV-2 S-binding antibodies. In b-d, each dot represents an individual sample.

Together, our results from four independent assays demonstrated the presence of pre-existing immunity to SARS-CoV-2 in uninfected individuals. Given their potential importance in shaping the natural course of SARS-CoV-2 infection, we next examined the prevalence of such cross-reactive antibodies in healthy donors. We further tested samples from a cohort of 50 SARS-CoV-2-uninfected pregnant women (Table S1), none of which had SARS-CoV-2 S-reactive antibodies of all three Ig classes (IgG, IgM and IgA) concurrently (Extended data Fig. 9). However, at least 5 of these samples (all collected in May 2018) showed evidence for lower levels of SARS-CoV-2 S-reactive IgG antibodies alone, which could not have been elicited by SARS-CoV-2 infection (Extended data Fig. 9). An additional 2 samples reacted with N, but not S or S1 on ELISA (Extended data Fig. 9). These data suggested that a minimum of 10% of this cohort had pre-existing antibodies cross-reactive with SARS-CoV-2 proteins. In a separate cohort of 101 SARS-CoV-2-uninfected donors (all sampled in May 2019), 3 had SARS-CoV-2 S-reactive IgG antibodies (Extended data Fig. 10), which did not correlate with antibodies to diverse viruses and bacteria also present in several of these samples (Table S1).

SARS-CoV-2 S-reactive IgM and IgA were also detected in 2 of these donors (aged 23 and 33), albeit at considerably lower levels than those in COVID-19 patients (Extended data Fig. 10), suggestive of recent or ongoing response.

The last two cohorts of SARS-CoV-2-uninfected individuals were sampled in late spring, a season when HCoV spread typically declines, and it was therefore less likely that they had experienced very recent HCoV infection. To examine if recent infection with a particular type of HCoV correlated with the presence of SARS-CoV-2 S-reactive antibodies, we tested an additional 13 donors who had each been infected with one of the 4 HCoV types between 12 and 55 days prior to sampling (Table S1). Of these 13 donors, only 1, recently infected with HCoV-OC43, had SARS-CoV-2 S-reactive IgG antibodies, and these at very low levels (Extended data Fig. 11), suggesting that their emergence is not simply a common transient event following each HCoV infection in this age group (median age 51 years; Table S1). Instead, given that HCoV-reactive antibodies are present in virtually all adults^34- 36^, the emergence of SARS-CoV-2 S-cross-reactive antibodies in a relatively small proportion of adults (15 of 262; 5.72%), indicates additional requirements, such as random B cell receptor repertoire focusing or context of initial exposure or frequency of HCoV infection, than time since the most recent HCoV infection. Indeed, the frequency of infection with each HCoV type displays a characteristic age distribution with children and adolescents experiencing the highest overall infection frequency^1,35-39^. We therefore examined a cohort of younger SARS-CoV-2-uninfected healthy donors (aged 1-16 years; Table S1). Remarkably, at least 21 of these 48 donors had readily detectable levels of SARS-CoV-2 S-reactive IgG antibodies (Fig. 4a,b). The prevalence of SARS-CoV-2 S-reactive IgG antibodies increased with age, peaking at 62% after 6 years of age (Fig. 4c), when HCoV seroconversion in this age group also peaks^34,35,37,38^, and was significantly higher than in adults (p<0.00001, Fisher’s exact test). Moreover, sera from SARS-CoV-2-uninfected children and adolescents efficiently neutralised SARS-CoV-2 S pseudotypes, but not VSVg pseudotypes *in vitro* (Fig. 4d). Sera from two SARS-CoV-2-uninfected adolescents, from which sufficient sample was available for testing, also neutralised authentic SARS-CoV-2 with comparable efficiency (Fig. 4e).

**Figure 4.**
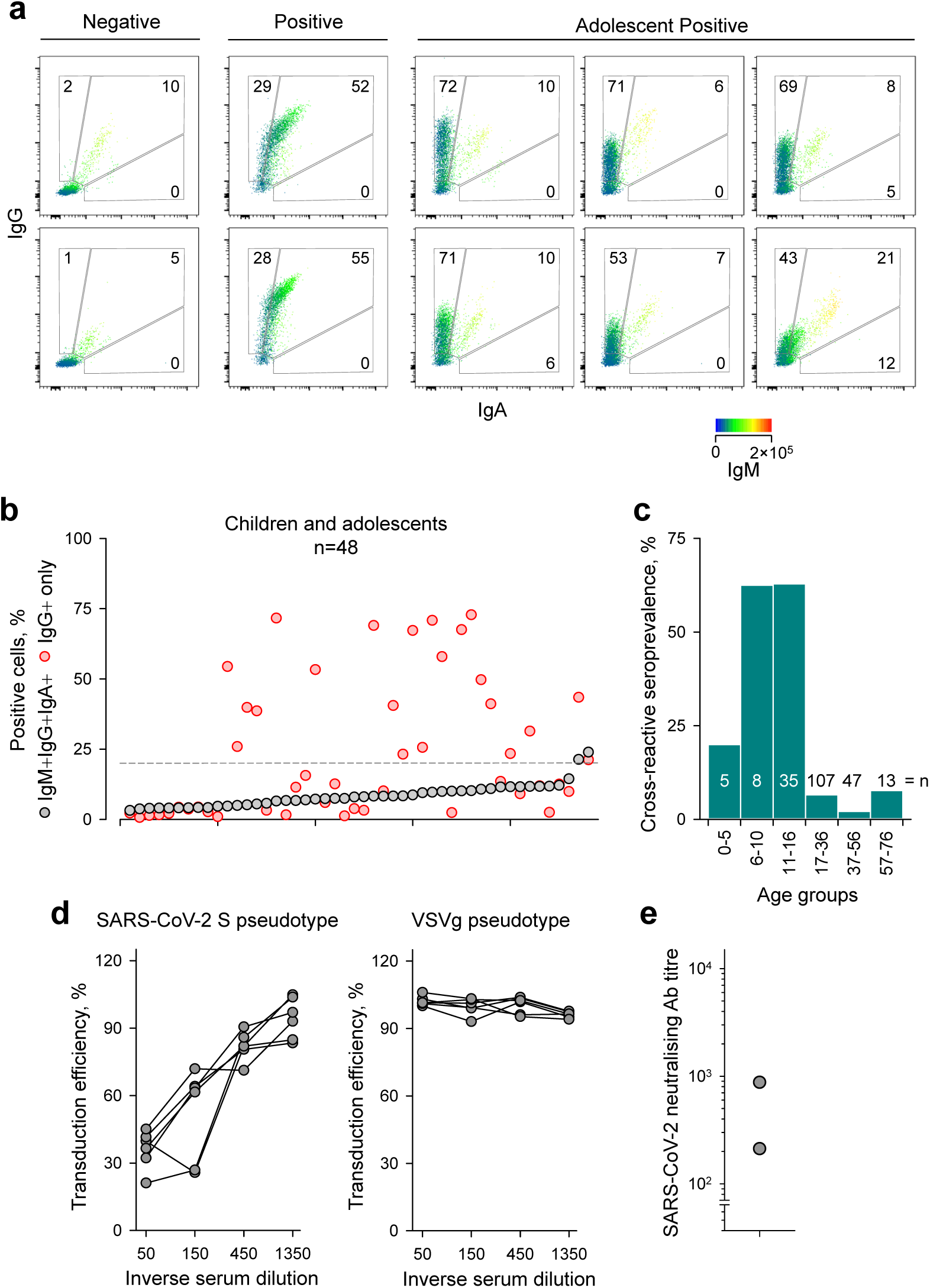
SARS-CoV-2-reactive antibody detection in children and adolescents. Samples from a total of 48 SARS-CoV-2-uninfected healthy children and adolescents collected between 2011 and 2018 were tested. **a**, Representative FACS profiles of control samples that were negative for all Ig classes (Negative) or COVID-19 patient samples (Positive), run in the same assay and of SARS-CoV-2-uninfected adolescents that were positive for SARS-CoV-2-cross-reactive antibodies. **b**, Frequency of cells that stained with all three antibody classes (IgM+IgG+IgA+) or only with IgG (IgG+) in each of these samples, ranked by their IgM+IgG+IgA+ frequency. **c**, Prevalence of SARS-CoV-2 S-cross-reactive antibodies in the indicated age groups. All samples used in this study for which the date of birth was known are included. **d**, Percent inhibition of transduction efficiency of SARS-CoV-2 S (*left*) or control VSVg (*right*) lentiviral pseudotypes by SARS-CoV-2-uninfected adolescent donor sera with FACS-detectable cross-relative antibodies. Each line is an individual serum sample. **e**, Authentic SARS-CoV-2 neutralisation titres of sera from two SARS-CoV-2-uninfected adolescents with FACS-detectable SARS-CoV-2 S-binding antibodies.

These observations support a model whereby exposure to HCoVs elicits humoral immunity that cross-reacts with conserved protein domains in other coronaviruses, including SARS-CoV-2. Infection with SARS-CoV-2 additionally induces *de novo* antibody responses to variable domains unique to this virus, specifically S1. Consistent with this model, SARS-CoV-2 S-reactive class-switched antibodies were detected by FACS concurrently with IgM antibodies in our COVID-19 patients, kinetics typically associated with anamnestic responses. A similar flow assay was recently used to demonstrate cross-reactivity of COVID-19 patient sera with the S proteins, but not the RBDs of the other two zoonotic CoVs, SARS-CoV and MERS^10^. Sera from SARS-CoV-recovered patients have been suggested to neutralise, to a degree, SARS-CoV-2 S pseudotypes^7^. Moreover, infection with SARS-CoV-2 has recently been shown to increase IgG seroreactivity to HCoVs^33^. SARS-CoV-2 S-reactive T helper cells, a prerequisite for class-switched antibody responses, have been detected in the majority of COVID-19 patients, as well as between one-third and two-thirds of healthy individuals, albeit at lower clonal frequencies^20,40^. In agreement with the specificity of the antibody response we describe here, T helper cells from COVID-19 patients targeted both subunits of SARS-CoV-2 S, whereas those from healthy individuals targeted the S2 subunit^20^.

Collectively, these studies highlight the existence of potentially functionally relevant antigenic epitopes in the S2 subunit, which is conserved among HCoVs and SARS-CoV-2. They echo findings with antigenic epitopes in the F subdomain of the influenza A virus hemagglutinin, which is also highly conserved among all influenza A virus 16 subtypes and the target of broadly neutralising antibodies^41,42^.

Over its entire length, SARS-CoV-2 S exhibits closer homology with the S proteins of the two betacoronaviruses, HCoV-OC43 and HCoV-HKU1, than of the alphacoronaviruses HCoV-NL63 and HCoV-229E (Extended data Fig. 12a). Infections with HCoV-OC43 are more prevalent than with HCoV-HKU1, also in the UK and in children and adolescents in particular^38^. Although these considerations would incriminate HCoV-OC43 as the likely inducer of SARS-CoV-2 S-reactive antibodies, other HCoVs are sufficiently homologous with SARS-CoV-2 to elicit cross-reactive humoral immune responses. To probe potential shared epitopes targeted in SARS-CoV-2 S, we constructed overlapping peptide arrays spanning the last 743 amino acids of SARS-CoV-2 S and tested the reactivity of adult and adolescent SARS-CoV-2-uninfected donor sera that had cross-reactive antibodies detectable by FACS (SARS-CoV-2 - Ab +) (Extended data Fig. 12b). As controls, we also used sera from seroconverted COVID-19 patients (SARS-CoV-2 + Ab +) and sera from adult SARS-CoV-2-uninfected donors without cross-reactive antibodies detectable by FACS (SARS-CoV-2 - Ab -). Multiple peptides in the array were recognised by the sera, identifying several putative epitopes in S2 (Fig. 5a,b; Extended data Fig. 12b). However, there were at least 5 epitopes recognised more strongly by sera with cross-reactive antibodies, particularly from adolescent donors, than by control sera (Fig. 5a,b). These included epitopes S_901-906_ (QMAYRF) and S_810-816_ (SKPSKRS) (Fig. 5), which were embedded in immunodominant regions previously experimentally defined in SARS-CoV and computationally predicted in SARS-CoV-2, respectively^43^. Both these epitopes were conserved between SARS-CoV and SARS-CoV-2, as well as among all four HCoVs (Fig. 5a,b). We further identified epitopes S_851-856_ (CAQKFN), S_1040-1044_ (VDFCG) and S_1205-1212_ (KYEQYIKW), all of which were highly conserved among HCoVs (Fig. 5a,b). Epitopes CAQKFN and KYEQYIKW mapped to a surface loop and the membrane-proximal region of S2, respectively, and were therefore missing from prior structures. However, the loop containing the CAQKFN epitope is present in the most recently reported structure^44^, where, similarly to other identified epitopes, it was located on the surface of S2 and accessible by antibodies (Fig. 5c). Lastly, epitope S_997-1002_ (ITGRLQ), also targeted more strongly by cross-reactive than control sera, was not located on the surface of pre-fusion S conformation (Fig. 5c), suggesting that additional epitopes might be accessible in alternative conformations, such as after the loss of the S1 subunit or after premature acquisition of the post-fusion conformation, as it was recently proposed^45,46^. Cross-reactivity with the identified epitopes was examined by ELISAs coated with synthetic peptides, which provided evidence for cross-reactivity (Extended data Fig. 13). Indeed, sera from seroconverted COVID-19 patients reacted with the selected peptides, as did those from SARS-CoV-2-uninfected adults or adolescents, in some cases more strongly than the former (Extended data Fig. 13). Reactivity with at least two peptides (containing epitopes CAQKFN and FIEDLLFN) appear restricted to SARS-CoV-2-uninfected donor samples with FACS-detectable cross-reactive antibodies and was not seen in control samples without FACS-detectable cross-reactive antibodies (Extended data Fig. 13).

**Figure 5.**
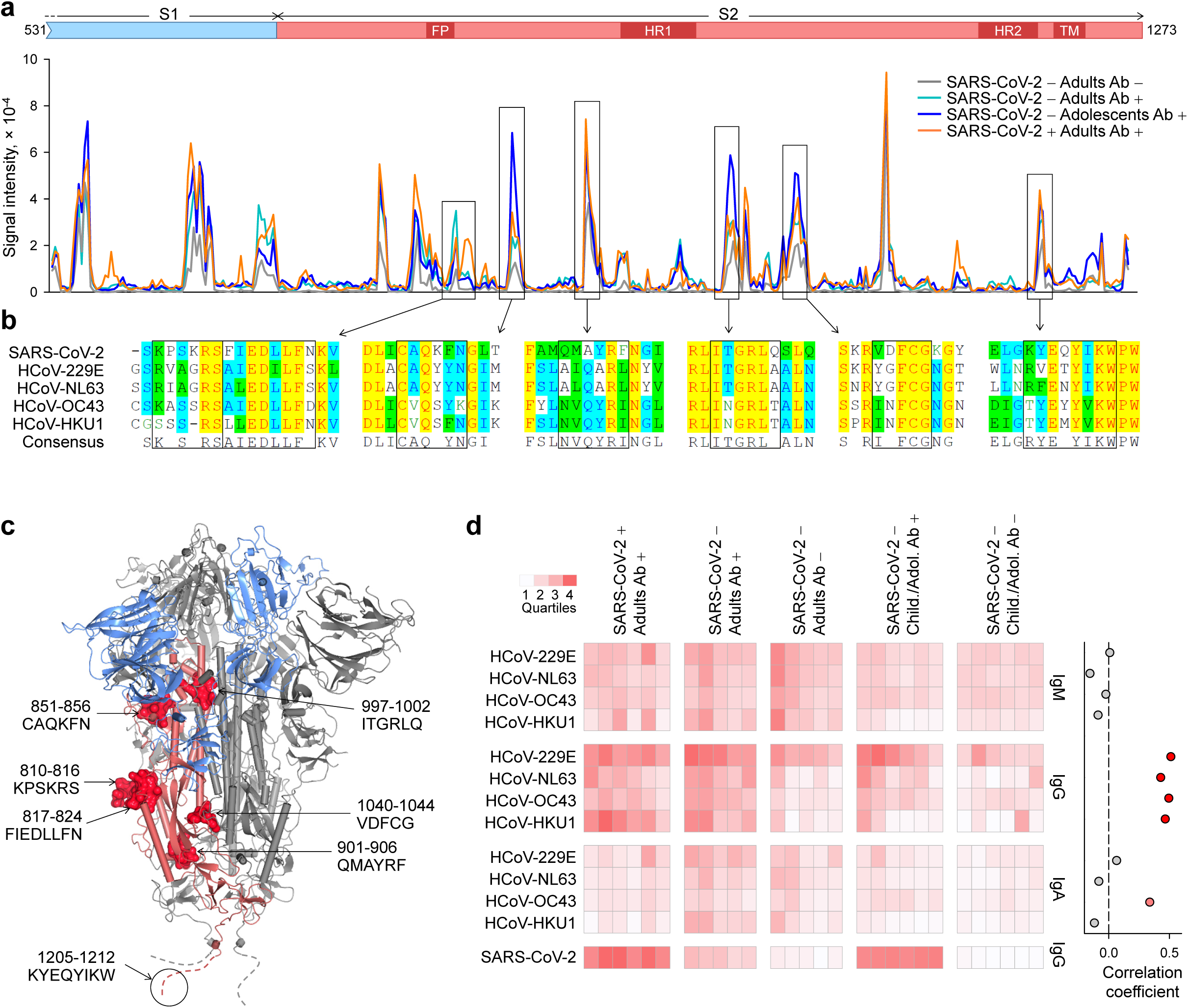
Mapping of cross-reactive epitopes in SARS-CoV-2 S. **a**, Schematic of the portion of SARS-CoV-2 S covered in the peptide arrays (*top*) and signal intensity obtained for each overlapping peptide along the length of the molecule (*bottom*), using pooled sera from the 4 indicated groups (1:100 final dilution). Boxed peaks indicate the regions that were recognised differentially by sera with and without SARS-CoV-2 S-cross-reactive antibodies, as determined by FACS. **b**, Alignment of the amino acid sequences of SARS-CoV-2 S and those of the 4 HCoV types. Similarity and homology is indicated with different shading and the consensus sequence is also shown. Boxes indicate the predicted core epitopes in each of the selected regions. **c**, Depiction of the linear predicted epitopes targeted by SARS-CoV-2 S-cross-reactive antibodies on the most complete structure of the trimeric SARS-CoV-2 spike (PDB ID: 6ZGE). One monomer is shown in colour, with S1 subunit coloured in blue and S2 in pale red, and the remaining two in grey. Peptides corresponding to the epitopes are shown for one monomer only; those present in the structure are depicted as crimson surfaces and the one derived from the membrane proximal region not present in the structure is represented as a circled dashed line. **d**, (*Left*), Percentage of cells transfected to express the S glycoproteins of each of the 4 HCoVs or of SARS-CoV-2 that were stained with the indicated sera. They include seroconverted adult COVID-19 patients (SARS-CoV-2 + Adults Ab +); SARS-CoV-2-uninfected adults or children/adolescents with (SARS-CoV-2 - Adults Ab + and SARS-CoV-2 - Child./Adol. Ab +, respectively) or without (SARS-CoV-2 - Adults Ab - and SARS-CoV-2 - Child./Adol. Ab -, respectively) FACS-detectable antibodies cross-reactive with SARS-CoV-2 S. Each column is an individual sample at 1:50 dilution, represented as a heatmap. Rows depict the staining for each antibody class. (Right), Coefficients of correlation between percentage of SARS-CoV-2 S-expressing cells stained with IgG antibodies and percentage of cells expressing the S glycoproteins of each of the 4 HCoVs stained with IgM, IgG or IgA antibodies.

These findings further supported the notion of pre-existing and *de novo* responses targeting conserved and unique SARS-CoV-2 S epitopes, respectively. However, the sequence conservation of the identified epitopes among the 4 HCoVs did not indicate if one was more likely than the others to have induced cross-reactive antibodies against these epitopes. We therefore examined reactivity of the sera against the S proteins of the 4 HCoVs separately using our FACS-based assay (Extended data Fig. 14). As expected, all sera exhibited some level of reactivity with one or more HCoVs, with HCoV-229E being the most frequent (Fig. 5d and Extended data Fig. 13). However, levels specifically of IgG antibodies against each of the 4 HCoV types, determined by this assay, were higher in sera from COVID-19 patients and SARS-CoV-2-infected donors with FACS-detectable cross-reactive antibodies to SARS-CoV-2 S, than the respective control sera (Fig. 5d). Indeed, IgG and IgA reactivity against HCoVs was higher in SARS-CoV-2-infected adults with, than without SARS-CoV-2-reactive IgG (p=1.4×10^−6^ for IgG and p=0.017 for IgA, Student’s t-test), and in SARS-CoV-2-infected children or adolescents with, than without SARS-CoV-2-reactive IgG (p=0.010 for IgG and p=0.021 for IgA, Student’s t-test), supporting a direct link between the two. Accordingly, IgG reactivity against each HCoV type was independently correlated with the presence of cross-reactive antibodies against SARS-CoV-2 (Fig. 5d). Weak correlation was also seen for IgA reactivity specifically against HCoV-OC43, but not the other HCoV or for IgM reactivity against any HCoV (Fig. 5d). Consistent with sequence conservation of the putative targeted epitopes, these data suggest the presence of IgG memory reactive with HCoVs, as well as SARS-CoV-2.

## Conclusions

Cross-reactivity between seasonal HCoVs and the pandemic SARS-CoV-2 needs to be carefully considered in the development and interpretation of assays for precise detection of SARS-CoV-2-specific antibodies. The FACS-based method we employed demonstrated the highest degree of specificity and sensitivity of the assays used here. This assay allows the distinction between pre-existing and *de novo* antibody responses to SARS-CoV-2, based on the levels of SARS-CoV-2 S-reactive IgG and parallel detection of IgM and IgA. These distinguishing features were maintained and, indeed, appear to increase during the first 43 days post onset of COVID-19 symptoms (the latest time-point we have examined). It is possible that these features are maintained for longer periods post SARS-CoV-2 infection, but it is also expected that the discriminating ability will be lost over time. Within each assay, increasing specificity by using higher positivity thresholds or omitting conserved protein domains may be at expense of sensitivity. This is evident in the use of less conserved SARS-CoV-2 protein domains such as S1 and RBD, which exhibited approximately 78% sensitivity during the first 43 days post onset of COVID-19 symptoms in our study, and which is likely to drop further with waning antibody titres over longer periods.

The ability of HCoV patient sera to neutralise authentic SARS-CoV-2 and SARS-CoV-2 S pseudotypes raise similar concerns regarding the specificity of neutralisation assays. For example, 5 of 1,000 samples from healthy Scottish blood donors collected in March 2020 neutralised SARS-CoV-2 S pseudotypes and 1 of 100 samples collected in 2019 also had neutralising activity in the absence of a strong ELISA signal^32^. Notably, these samples were described to have low or no SARS-CoV-2 S-reactive IgM antibodies^32^, a feature we would associate with immune memory of HCoVs.

In addition to its implications for serology assay development and interpretation or for the design of vaccination studies, potential cross-reactivity between seasonal HCoVs and the pandemic SARS-CoV-2 has important ramifications for natural infection. Thorough epidemiological studies of HCoV transmission suggest that cross-protective immunity is unlikely to be sterilising or long-lasting^39^, which is also supported by repeated reinfection of all age groups^4^, sometimes even with homologous HCoVs^47^. Nevertheless, prior immunity induced by one HCoV has also been reported to reduce the transmission of homologous and, importantly, heterologous HCoVs, and to ameliorate the symptoms where transmission is not prevented^1,4,5^. A possible modification of COVID-19 severity by prior HCoV infection might account for the age distribution of COVID-19 susceptibility, where higher HCoV infection rates in children than in adults^5,35,37^, correlates with relative protection from COVID-19^48^, and might also shape seasonal and geographical patterns of transmission.

Public health measures intended to prevent the spread of SARS-CoV-2 will also prevent the spread of and, consequently, maintenance of herd immunity to HCoVs, particularly in children. It is, therefore, imperative that any effect, positive or negative, of pre-existing HCoV-elicited immunity on the natural course of SARS-CoV-2 infection is fully delineated.

## Methods

### Patients and clinical samples

Serum or plasma samples were obtained from University College London Hospitals (UCLH) COVID-19 patients testing positive for SARS-CoV-2 infection by RT-qPCR and sampled between March 2020 and April 2020. An initial cohort of 35 patients (31 annotated) and an extended cohort of 135 patients were tested between 2 and 43 days after the onset of COVID-19 symptoms (Table S1). A total of 262 samples form SARS-CoV-2-uninfected adults and 48 SARS-CoV-2-uninfected children and adolescents were also used (described in Table S1). Samples from adults were obtained from UCLH and Public Health Wales, University Hospital of Wales, and samples from children and adolescents were obtained from the Centre for Adolescent Rheumatology Versus Arthritis at University College London (UCL), UCLH, and Great Ormond Street Hospitals (GOSH), and Great Ormond Street Institute for Child Health (ICH) with ethical approval (refs 11/LO/0330 and 95RU04). All patient sera and sera remaining after antenatal screening of healthy pregnant women were from residual samples prior to discarding, in accordance with Royal College Pathologists guidelines and the UCLH Clinical Governance for assay development and GOSH and ICH regulations. All serum or plasma samples were heat-treated at 56°C for 30 min prior to testing.

### Viral infection RT-qPCR diagnosis

SARS-CoV-2 nucleic acids were detected at Health Services Laboratories, London, UK, by an RT-qPCR method as recently described^49^. HCoV nucleic acids were detected by RT-qPCR, as part of a diagnostic panel for respiratory viruses, run by Health Services Laboratories (HSL), London, UK.

### Cell lines and virus

HEK293T cells were obtained from the Cell Services facility at The Francis Crick Institute, verified as mycoplasma-free and validated by DNA fingerprinting. Vero-E6 cells were from the National Institute for Biological Standards and Control, UK. HEK293T cells overexpressing ACE2 were generated by transfection, using GeneJuice (EMD Millipore), with a plasmid containing the complete human ACE2 transcript variant 1 cDNA sequence (NM_001371415.1) cloned into the mammalian expression vector pcDNA3.1-C’ FLAG by Genscript. Cells were grown in Iscove’s Modified Dulbecco’s Medium (Sigma Aldrich) supplemented with 5% fetal bovine serum (Thermo Fisher Scientific), L-glutamine (2 mmol/L, Thermo Fisher Scientific), penicillin (100 U/ml, Thermo Fisher Scientific), and streptomycin (0.1 mg/ml, Thermo Fisher Scientific). The SARS-CoV-2 isolate hCoV-19/England/02/2020 was obtained from the Respiratory Virus Unit, Public Health England, UK, (GISAID EpiCov^™^ accession EPI_ISL_407073) and propagated in Vero E6 cells.

### Flow cytometry

HEK293T cells were transfected with an expression vector (pcDNA3) carrying a codon-optimised gene encoding the wild-type SARS-CoV-2 S (UniProt ID: P0DTC2) (kindly provided by Massimo Pizzato, University of Trento, Italy), using GeneJuice (EMD Millipore). Similarly, HEK293T cells were transfected with expression vectors (pCMV3) expressing HCoV-229E S (UniProt ID: APT69883.1), HCoV-NL63 S (UniProt ID: APF29071.1), HCoV-OC43 S (UniProt ID: AVR40344.1) or HCoV-HKU1 S (UniProt ID: Q0ZME7.1) (all from SinoBiological). Two days after transfection, cells were trypsinised and transferred into V-bottom 96-well plates (20,000 cells/well). Cells were incubated with sera (diluted 1:50 in PBS) for 30 min, washed with FACS buffer (PBS, 5% BSA, 0.05% Tween 20, 0.05% sodium azide), and stained with BV421 anti-IgG (Biolegend), APC anti-IgM (Biolegend) and PE anti-IgA (Miltenyi Biotech) for 30 min (all antibodies diluted 1:200 in FACS buffer). Cells were washed with FACS buffer and fixed for 20 min in CellFIX buffer (BD Bioscience). Samples were run on a Ze5 (Bio-Rad) running Bio-Rad Everest software v2.4 or an LSR Fortessa with a high-throughput sampler (BD Biosciences) running BD FACSDiva software v8.0, and analysed using FlowJo v10 (Tree Star Inc.) analysis software. Transfection efficiencies were determined by staining with a fixed concentration of the S1-reactive CR3022 antibody (100 ng/ml) (Absolute Antibodies) and control convalescent sera (1:50 dilution), followed by BV421 anti-IgG antibody. Staining with convalescent sera, detecting epitopes over the entire S ectodomain, was consistently higher than staining with CR3022, detecting only the S1 epitope, and was taken as the maximum transfection efficiency. This varied between 68% and 95% between experiments.

### Recombinant protein production

The SARS-CoV-2 RBD and S1 constructs, spanning SARS-CoV-2 S (UniProt ID: P0DTC2) residues 319-541 (RVQPT…KCVNF) and 1-530 (MFVFL…GPKKS), respectively, were produced with C-terminal twin Strep tags. To this end, the corresponding codon-optimised DNA fragments were cloned into mammalian expression vector pQ-3C-2xStrep^50^. A signal peptide from immunoglobulin kappa gene product (METDTLLLWVLLLWVPGSTGD) was used to direct secretion of the RBD construct. Stabilised ectodomain of the SARS-CoV-2 S glycoprotein (residues 1-1208) with inactivated furin cleavage site (RRAR, residues 682-685 mutated to GSAS) and a double proline substitution (K986P/V987P)^51,52^ was produced with a C-terminal T4 fibritin trimerization domain and a hexahistidine (His_6_) tag from the pcDNA3 vector. Expi293F cells growing at 37^°^C in 5% CO_2_ atmosphere in shake flasks in FreeStyle 293 medium were transfected with the corresponding plasmids using ExpiFectamine reagent (Thermo Fisher Scientific). Conditioned medium containing secreted proteins was harvested twice, 3-4 and 6-8 days post-transfection. Twin Strep- and His_6_-tagged proteins were captured on Streptactin XT (IBA LifeSciences) or Talon (Takara) affinity resin, respectively, and purified to homogeneity by size exclusion chromatography through Superdex 200 (GE Healthcare) in 150 mM NaCl, 1 mM EDTA, 20 mM Tris-HCl, pH 8.0. Full-length SARS CoV2 N gene product was produced with an N-terminal His_6_ tag from pOPTH-1124 plasmid (kindly provided by Jakub Luptak and Leo James, Laboratory for Molecular Biology, Cambridge, UK). *Escherichia coli* C43(DE3) cells (Lucigen) transformed with pOPTH-1124 were grown in terrific broth medium, and expression was induced by addition of 1 mM Isopropyl β-D-1-thiogalactopyranoside at 37^°^C. Bacteria, harvested 4 h post-induction, were disrupted by sonication in core buffer (1M NaCl, 10 mM imidazole, 20 mM HEPES-NaOH, pH 8.0) supplemented with BaseMuncher nuclease (Expedion; 1 ml per 40 ml cell suspension) and Complete EDTA-free protease inhibitor mix (Roche). The extract was precleared by centrifugation, and His_6_-tagged protein was captured on NiNTA agarose (Qiagen). Following extensive washes with core buffer supplemented with 20 mM imidazole, the protein was eluted with 500 mM imidazole. SARS CoV-2 N, was further purified by cation exchange and heparin affinity chromatography prior polishing by gel filtration through a Superdex 200 16/40 column (GE Healthcare), which was operated in 300 mM NaCl, 20 mM HEPES-NaOH, pH 8.0. Purified SARS CoV2 antigens, concentrated to 1-5 mg/ml by ultrafiltration using appropriate VivaSpin devices (Sartorius), were snap-frozen in liquid nitrogen in small aliquots and stored at -80^°^C.

### ELISA

96-well MaxiSorp plates (Thermo Fisher Scientific) were coated overnight at 4^°^C with purified protein in PBS (3 µg/ml per well in 50 µL) and blocked for 1 hr in blocking buffer (PBS, 5% milk, 0.05% Tween 20, 0.01% sodium azide). Sera was diluted in blocking buffer (1:50) and 50 µL added to the plate then incubated for 1 hr at room temperature. After washing 4 times with PBS-T (PBS, 0.05% Tween 20), plates were incubated with alkaline phosphatase-conjugated goat anti-human IgG (1:1000, Jackson ImmunoResearch) for 1 hr. Plates were developed by adding 50 µL of alkaline phosphatase substrate (Sigma Aldrich) for 15-30 min after 6 washes with PBS-T. Optical densities were measured at 405 nm on a microplate reader (Tecan). CR3022 (Absolute Antibodies) was used as a positive control for ELISAs coated with S, S1, and RBD. For ELISAs with synthetic peptides, 96-well Nunc Immobilizer Amino Plates (Thermo Fisher Scientific) were used for coating overnight at 4^°^C with peptides (10 µg/ml per well in 50 µL)

### Lentiviral particle production and neutralisation

Lentiviral particles pseudotyped with either SARS-CoV-2 S or Vesicular Stomatitis Virus glycoprotein (VSVg) were produced by co-transfection of HEK293T cells with plasmids encoding either of these glycoproteins together with a plasmid encoding the SIVmac Gag-Pol polyprotein and a plasmid expressing an HIV-2 backbone with a GFP encoding gene, using GeneJuice (EMD Millipore). Virus-containing supernatants were collected 48 hr post-transfection and stored at -80^°^C until further use. For neutralisation assays, lentiviral pseudotypes were incubated with serial dilutions of patient sera at 37^°^C for 30 minutes and were subsequently added to HEK293T cells seeded in 96-well plates (3,000 cells/well). Polybrene (4 ug/mL, Sigma Aldrich) was also added to the cells and plates were spun at 1,200 rpm for 45 min. The percentage of transduced (GFP+) cells was assessed by flow cytometry 72 hours later. The inverse serum dilution leading to 50% reduction of GFP+ cells was taken as the neutralising titre.

### SARS-CoV-2 plaque reduction neutralisation test

Confluent monolayers of Vero E6 cells were incubated with 10-20 plaque-forming units (PFU) of SARS-CoV-2 strain hCoV-19/England/2/2020 and two-fold serial dilutions of human sera (previously heat-treated at 56°C for 30 min) starting at 1:40 dilution, for 3 hours at 37°C, 5% CO_2_, in triplicate per condition. The inoculum was then removed and cells were overlaid with virus growth medium containing avicel at a final concentration of 1.2%. Cells were incubated at 37°C, 5% CO_2_. At 24 hours post-infection, cells were fixed in 4% paraformaldehyde and permeabilised with 0.2% Triton-X-100/PBS and virus plaques were visualised by immunostaining, as described previously for the neutralisation of influenza viruses^53^, except using a rabbit polyclonal anti-NSP8 antibody and anti-rabbit-HRP conjugate and detected by action of HRP on a tetra methyl benzidine (TMB) based substrate. Virus plaques were quantified and IC_50_ for sera was calculated using LabView software as described previously^53^.

### Peptide arrays

Peptide arrays were synthesised on an Intavis ResPepSL Automated Peptide Synthesiser (Intavis Bioanalytical Instruments, Germany) on a cellulose membrane by cycles of N(a)-Fmoc amino acids coupling via activation of carboxylic acid groups with diisopropylcarbodiimide (DIC) in the presence of Ethyl cyano(hydroxyimino)acetate (Oxyma pure) followed by removal of the temporary α-amino protecting group by piperidine treatment. Subsequent to chain assembly, side chain protection groups were removed by treatment of membranes with a deprotection cocktail (20 mls 95% trifluoroacetic acid, 3% triisopropylsilane 2% water 4 hours at room temperature) then washing (4 × DCM, 4 × EtOH 2 × H20, 1 × EtOH) prior to being air dried. Membranes were blocked for 1 hour in blocking buffer (PBS, 5% BSA, 0.05% Tween 20, 0.05% sodium azide), then stained with pooled sera (1:100 in blocking buffer) for 2 hours at room temperature. Membranes were washed 3 times in PBS-T, then stained with IRDye 800CW Goat anti-Human IgG (Licor; 1:15,000 in blocking buffer) for 1 hour at room temperature in the dark. Membranes were washed 3 times in PBS-T and once in PBS before imaging on an Odyssey CLx Infrared scanner (Licor). Scanned images were analysed in Image Studio v5.2 (Licor).

### Individual peptide synthesis

Individual peptides were synthesised on an Intavis ResPepSLi Automated Peptide Synthesiser (Intavis Bioanalytical Instruments, Germany) on Rink amide resin (0.26 mmole/g, 0.1 mmol) using N(a)-Fmoc amino acids and HATU as the coupling reagent. Following amino acid chain assembly, peptides were cleaved from the resin by addition to cleavage cocktail (10 ml, 92.5% TFA, 2.5% H2O, 2.5% EDT, 2.5% TIS) for 2 hours. Following resin removal, peptide precipitation and extensive washing with ether, the peptides were analysed by LC–MS on an Agilent 1100 LC-MSD.

### Data analysis

Data were analysed and plotted in GraphPad Prism v8 (GraphPad Software) or SigmaPlot v14.0 (Systat Software). Sequence alignments were performed with Vector NTI v11.5 (Thermo Fisher Scientific).

## Acknowledgements

We thank L. James and J. Luptak (Laboratory for Molecular Biology, Cambridge, UK) for a generous gift of SARV CoV2 N expression construct and M. Pizzato (University of Trento, Italy) for providing codon-optimised SARS CoV2 S cDNA. We also thank the entire CRICK COVID-19 Consortium. We are grateful for assistance from the Cell Services and High Throughput Screening facilities at the Francis Crick Institute and UCLH Biochemistry (A. Goyale and C. Wilson) and to Mr Michael Bennet and Mr Simon Caidan for training and support in the high-containment laboratory. This work was supported by a Centre of Excellence Centre for Adolescent Rheumatology Versus Arthritis grant, 21593, as well as support from the Great Ormond Street Childrens Charity, CureJM Foundation and the NIHR Biomedical Research Centres at GOSH and UCLH. This work was supported by the Francis Crick Institute, which receives its core funding from Cancer Research UK, the UK Medical Research Council, and the Wellcome Trust.

---

**Table S1.** Details of patient and healthy donor samples used in this study. Samples were from healthy volunteers at the UCL Centre for Adolescent Rheumatology, Great Ormond Street (GOSH) Institute for Child Health (ICH) and Adolescent Centre Biobank.

| Number of donors | Median age (range) | Date of blood collection | Median number of days (range) since |  |
| --- | --- | --- | --- | --- |
|  |  |  | symptoms onset | RT-qPCR confirmation |
| **A. Initial cohort of COVID-19 patients** |  |  |  |  |
| 35 | 64 (24 – 90) | 3/2020 – 4/2020 | 14 (5 – 29) | 8 (-3 – 17) |
| Patients at UCLH. Age was available for 31 patients. |  |  |  |  |
| **B. Extended cohort of COVID-19 patients** |  |  |  |  |
| 135 | 64 (36 – 90) | 4/2020 | 20 (2 – 43) | 13 (-1 – 33) |
| Patients at UCLH. Age was available for 80 patients. |  |  |  |  |
| **C. SARS-CoV-2-uninfected patients without recent HCoV infection (SARS-CoV-2 - HCoV -)** |  |  |  |  |
| 31 | n/a | 8/2019 – 9/2019 | n/a | n/a |
| Haematology patients at UCLH testing negative for HCoV infection. |  |  |  |  |
| **D. SARS-CoV-2-uninfected patients with recent HCoV infection (SARS-CoV-2 - HCoV +)** |  |  |  |  |
| 34 | n/a | 12/2019 – 3/2020 | n/a | 18 (-5 – 119) |
| Haematology patients at UCLH testing positive for HCoV infection. |  |  |  |  |
| **E. SARS-CoV-2-uninfected patients of unknown recent HCoV status (SARS-CoV-2 -)** |  |  |  |  |
| 30 | n/a | 8/2019 – 9/2019 | n/a | n/a |
| Patients at UCLH, not tested for HCoV infection. |  |  |  |  |
| **F. SARS-CoV-2-uninfected pregnant women** |  |  |  |  |
| 50 | 32 (17 – 52) | 5/2018 | n/a | n/a |
| Healthy visitors of antenatal clinics at UCLH. |  |  |  |  |
| **G. SARS-CoV-2-uninfected patients with unrelated infections** |  |  |  |  |
| 101 | 31 (18 – 65) | 5/2019 | n/a | n/a |
| Patients tested at UCLH. They include patients testing positive for antibodies to Influenza HA (2), HBV S (13), HBV C (4), HAV (1), EBV (1), VZV (1), Borrelia sp. (Lyme disease) (1) and Treponema sp. (syphilis) (1). |  |  |  |  |
| **H. SARS-CoV-2-uninfected patients with recent HCoV infection of known type** |  |  |  |  |
| 13 | 52 (21 – 75) | 1/2019 – 4/2020 | n/a | 18 (12 – 55) |
| Patients at University Hospital of Wales. A total of 16 samples were taken from these 13 patients. One haematology patient, persistently infected with NL63 was sampled 4 separate times and all other donors were sampled once. They include patients infected with OC43 (5), NL63 (3), 229E (2) and HKU1 (3). |  |  |  |  |
| **I. SARS-CoV-2-uninfected Healthy Children and Adolescents** |  |  |  |  |
| 48 | 14 (1 – 16) | 4/2011 – 12/2018 | n/a | n/a |
| Samples were from healthy volunteers at the UCL Centre for Adolescent Rheumatology, Great Ormond Street (GOSH) Institute for Child Health (ICH) and Adolescent Centre Biobank. |  |  |  |  |

**Extended data Fig. 1.**
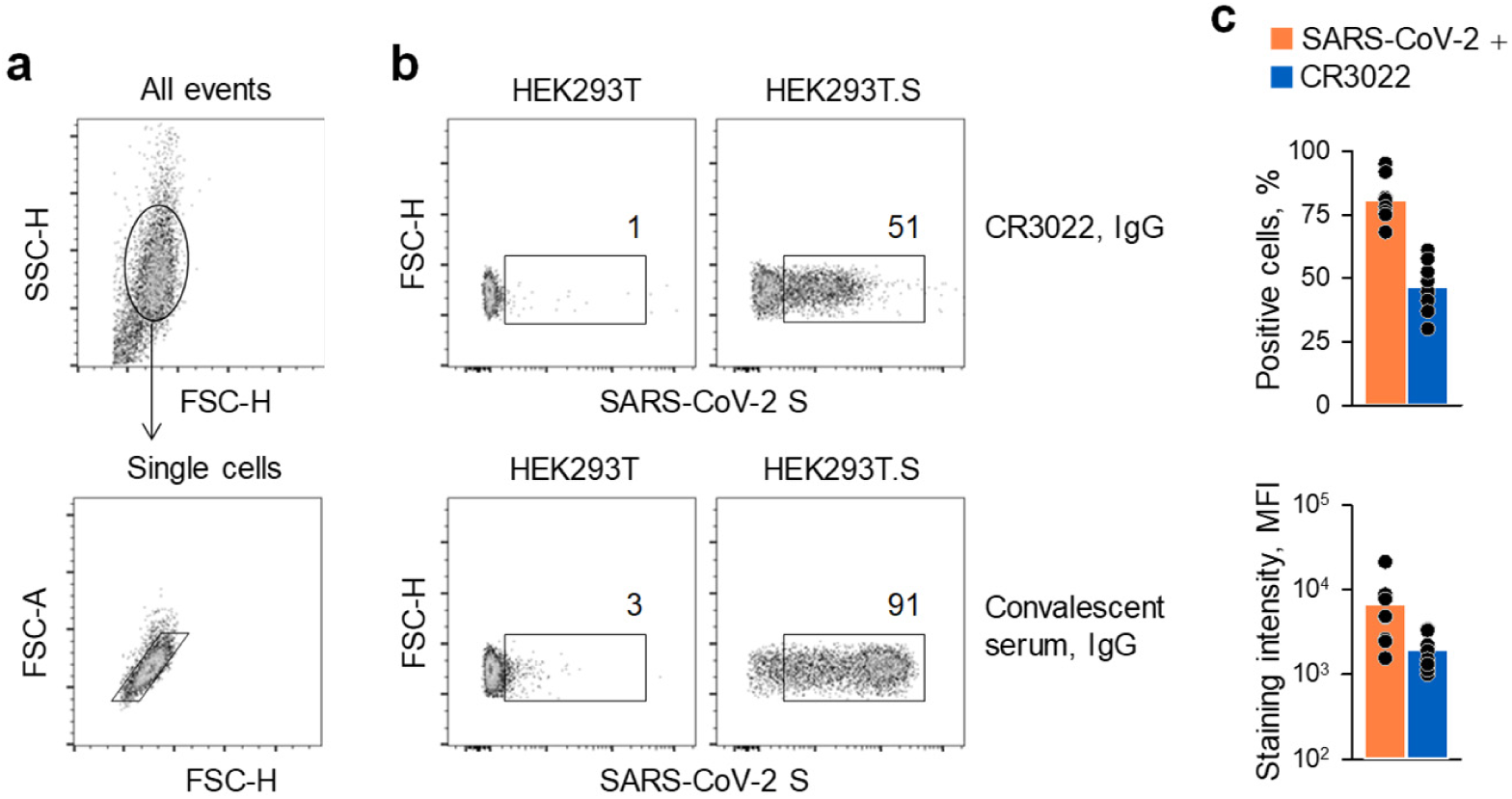
Detection of cell membrane-bound SARS-CoV-2 S by FACS. HEK293T cells were transfected with an expression plasmid encoding the wild-type SARS-CoV-2 S and two days later were used for FACS analysis. **a**, Gating of HEK293T cells and of single cells in cell suspensions. **b**, Representative staining of untransfected HEK293T cells or SARS-CoV-2 S-transfected HEK293T cells (HEK293T.S) with either the CR3022 monoclonal antibody or serum from COVID-19 patients as the primary antibody, followed by an anti-human IgG secondary antibody. **c**, Percentage of positive cells (*top*) and mean fluorescence intensity (MFI) (*bottom*) of staining with either the CR3022 antibody or COVID-19 patient serum in 8 independent experiments. The differences in the percentage of positive cells and MFI were statistically significant with p=0.000008, paired t-test and p=0.008, Wilcoxon Signed Rank Test, respectively.

**Extended data Fig. 2.**
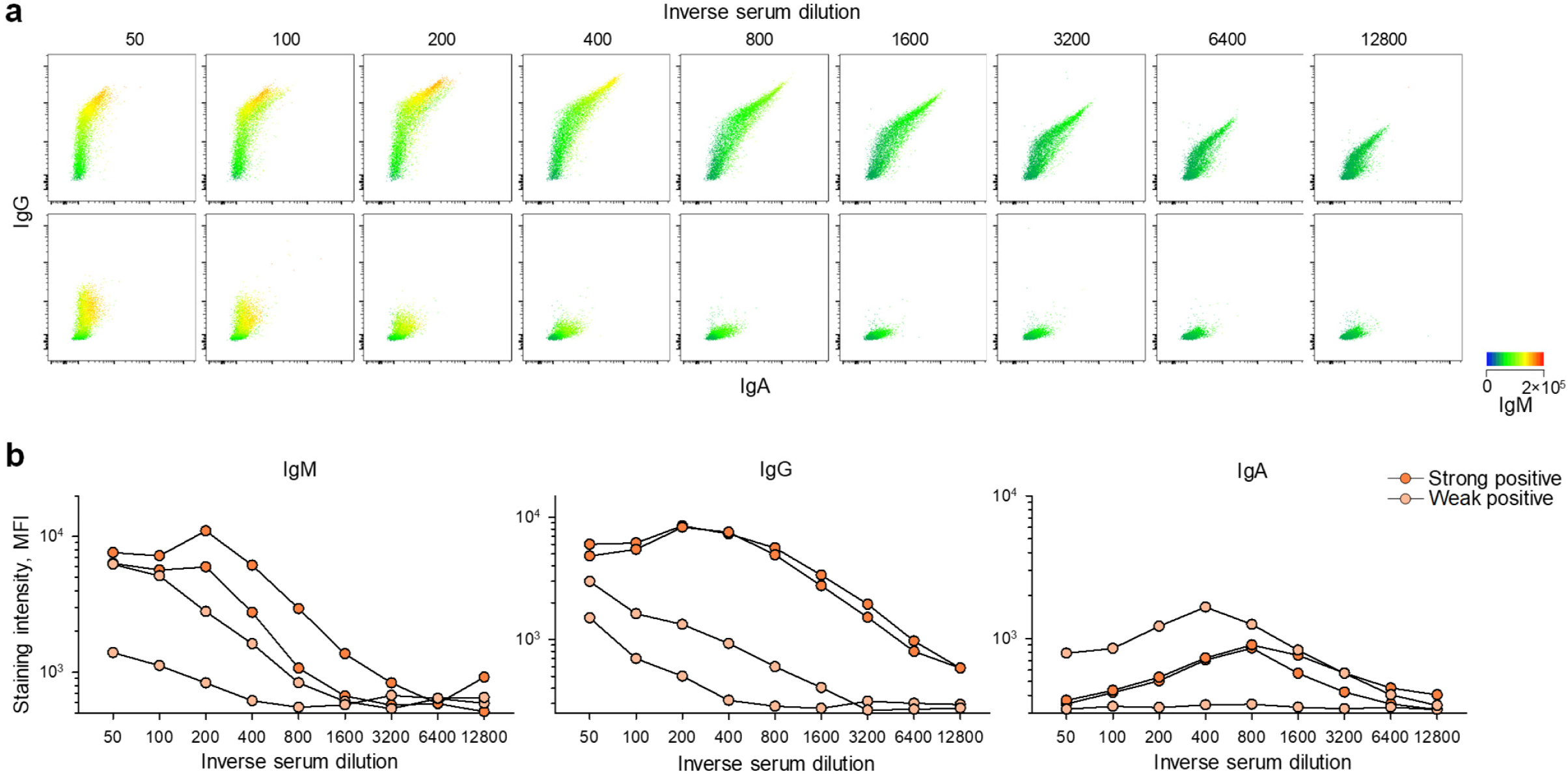
Detection of SARS-CoV-2 S-binding IgM, IgG and IgA antibodies by FACS. **a**, FACS profiles of SARS-CoV-2 S-transfected HEK293T cells staining with the indicated serial dilutions of COVID-19 patient sera selected to represent the high (*top*) and low (*bottom*) ends of detection at 1:50 dilution. Levels of IgA and IgG are indicated in the x and y axes, respectively, and levels of IgM are indicated by a heatmap. **b**, Mean fluorescence intensity (MFI) of IgM, IgG and IgA staining, according to the indicated serial dilution of two samples from each end of the detection range. Each line represents an individual COVID-19 patient serum sample.

**Extended data Fig. 3.**
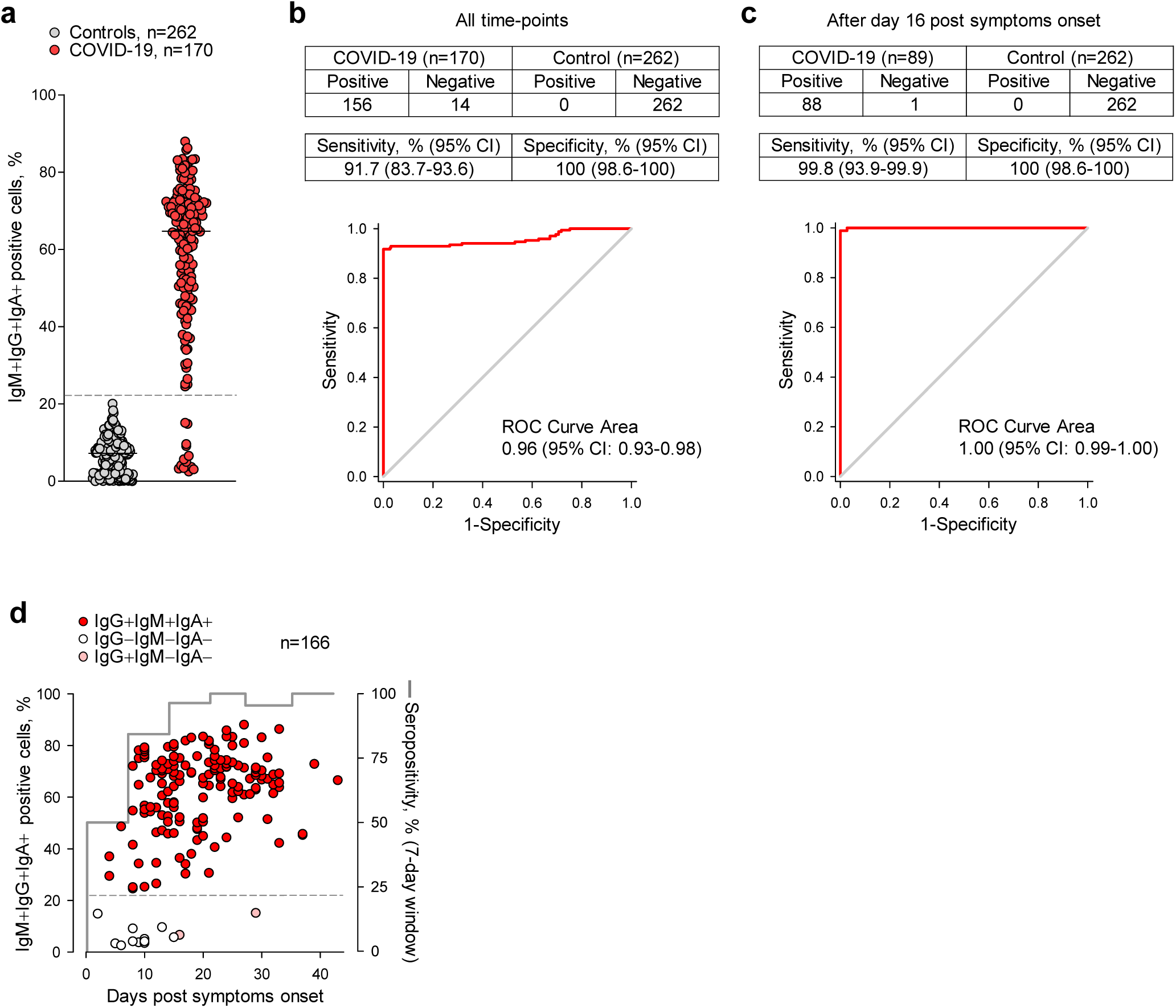
Performance of the FACS-based assay for SARS-CoV-2-reactive antibody detection. **a**, Frequency of SARS-CoV-2 S-transfected HEK293T cells that stained positive with all three antibody classes (IgM+IgG+IgA+) with sera from 170 confirmed COVID-19 cases and 262 controls. The dashed line represents the calculate cut-off for positivity. **b-c**, Sensitivity and specificity for the assay calculated for all the sample collected at any time-point post symptoms onset (b) or only for samples collected 16 days after the onset of COVID-19 symptoms (c). The receiver operating characteristics (ROC) curve areas and 95% confidence intervals (CI) are also shown for each group. **d**, Kinetics of seroconversion in COVID-19 patients determined by the FACS-based assay. Percentages of IgM+IgG+IgA+ positive cells are plotted over time of sample collection since symptoms onset. Seropositivity was calculated for each consecutive week since symptoms onset. Only COVID-19 patients with known date of symptoms onset are included.

**Extended data Fig. 4.**
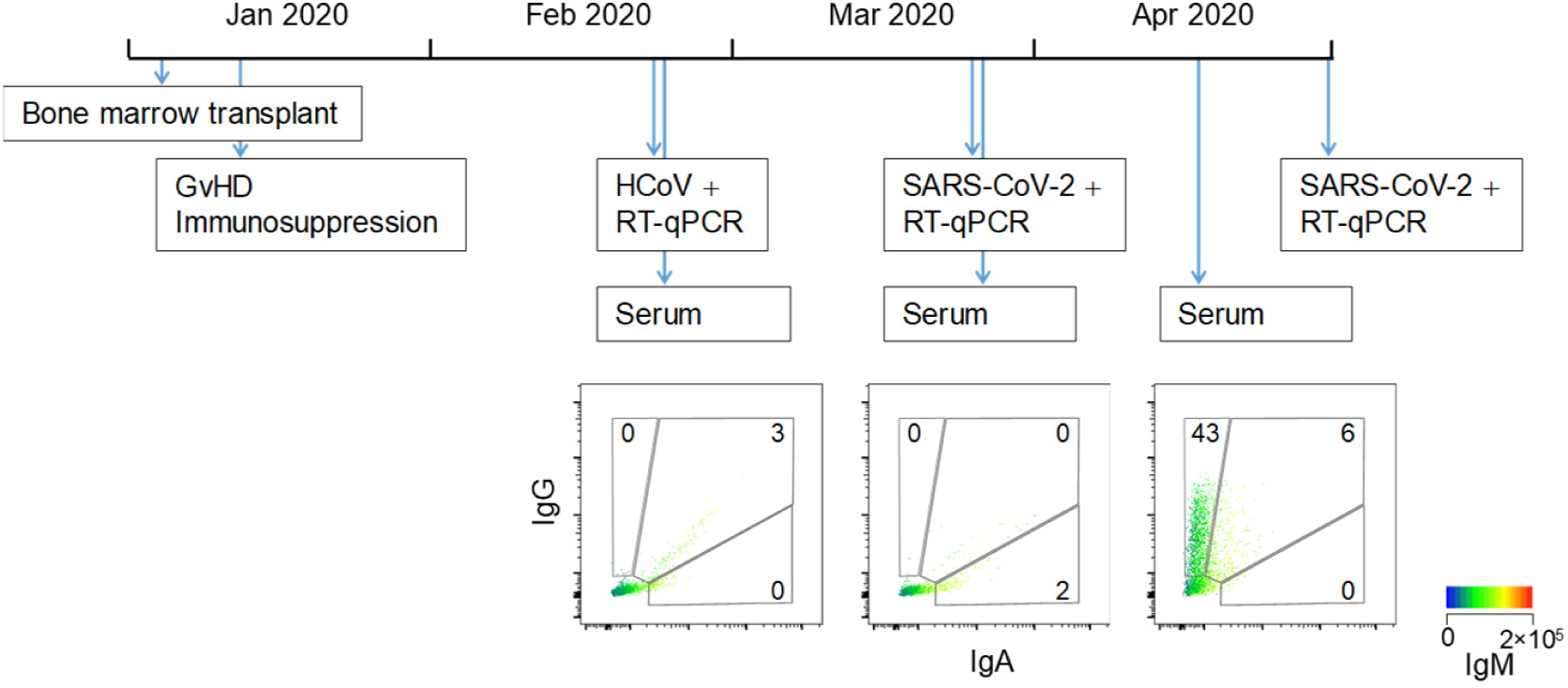
Sequential HCoV and SARS-CoV-2 infection and clinical observations in an individual case. A bone marrow transplant recipient, with graft versus host disease (GvHD) and associated immunosuppression tested positive by RT-qPCR for HCoV in February 2020. The patient’s sera did not contain SARS-CoV-2 S-binding antibodies at that time. The patient later acquired likely nosocomial SARS-CoV-2 infection with first RT-qPCR confirmation in March 2020. The patient’s sera remained negative for SARS-CoV-2 S-binding antibodies at the second sampling time-point, but exhibited an only IgG+ profile three weeks later (April 2020). The patient experienced mild COVID-19 symptoms that did not require hospitalisation, but remained positive for SARS-CoV-2 infection, with RT-qPCR confirmation in late April 2020. None of the serial serum samples had significant neutralising activity against SARS-CoV-2 pseudotypes.

**Extended data Fig. 5.**
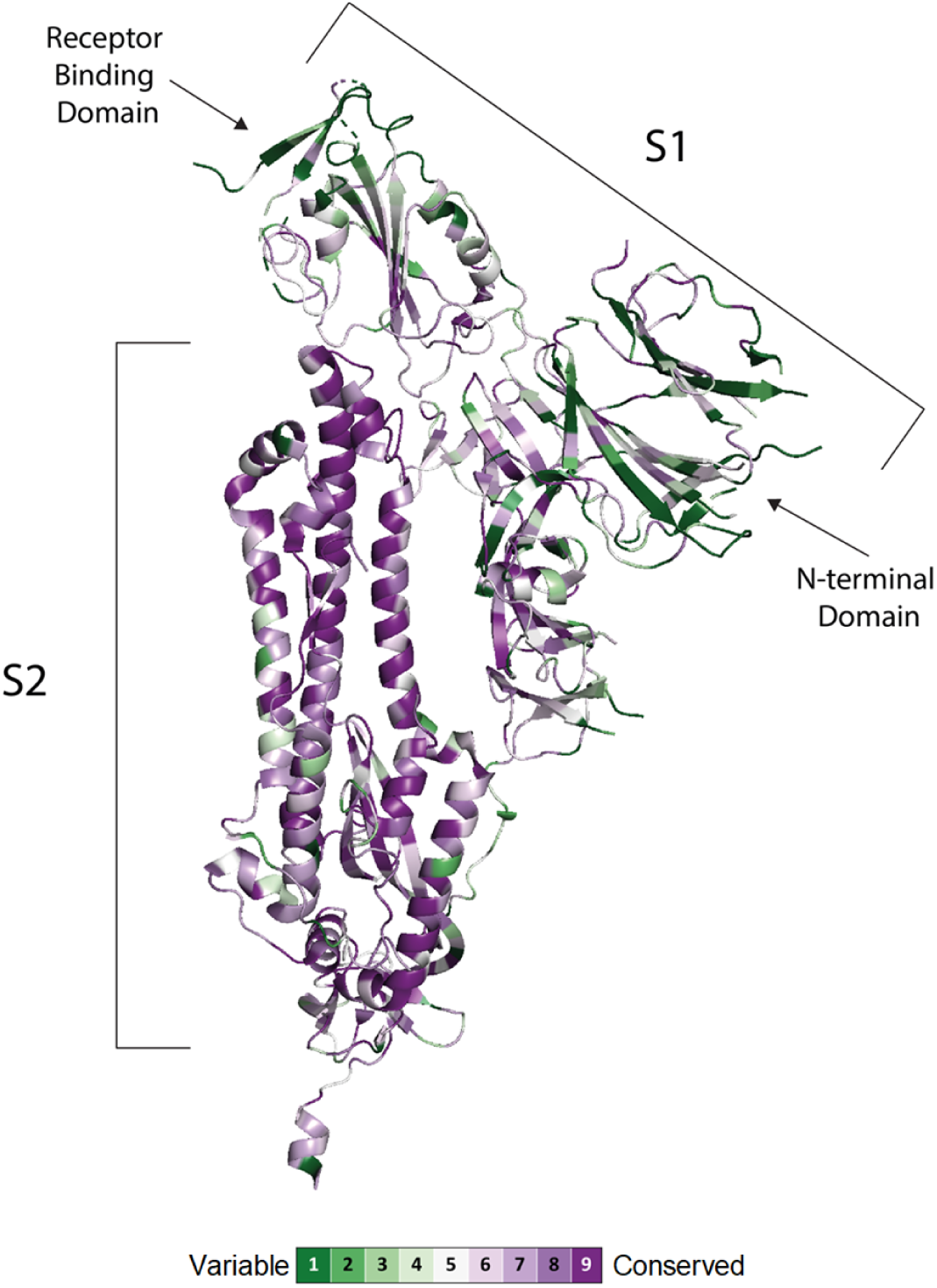
Conservation of SARS-CoV-2 S subunits. Structure of the SARS-CoV-2 S protomer with each amino acid residue coloured according to conservation among 24 animal and human CoVs. The alignment and the figure were generated using the ConSurf algorithm (https://consurf.tau.ac.il) using a single chain of the SARS-CoV-2 spike protein in the closed state (PDB ID 6VXX) as a reference.

**Extended data Fig.6.**
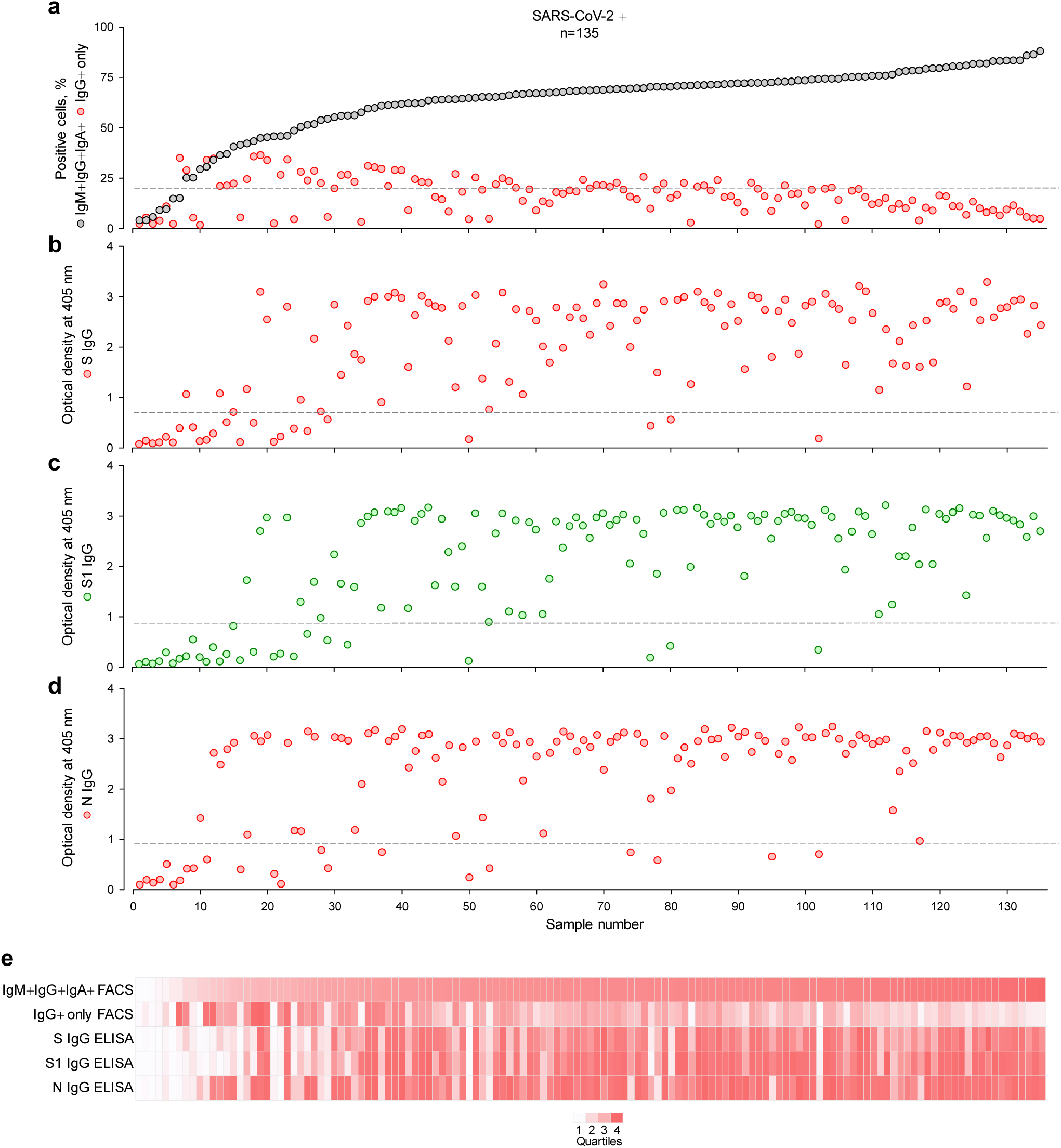
Comparison of antibody detection methods in an extended cohort of COVID-19 patients. Samples from a total of 135 confirmed COVID-19 patients were tested (Table S1). **a**, Frequency of cells that stained with all three antibody classes (IgM+IgG+IgA+) or only with IgG (IgG+) in each of these samples, ranked by their IgM+IgG+IgA+ frequency. **b-d**, Optical densities from ELISAs coated with S (b), S1 or RBD (c) or N (d) of the same samples. Dashed lines in a-d denote the assay sensitivity cut-offs. **e**, Summary of the results from a-d, represented as a heatmap of the quartile values.

**Extended data Fig. 7.**
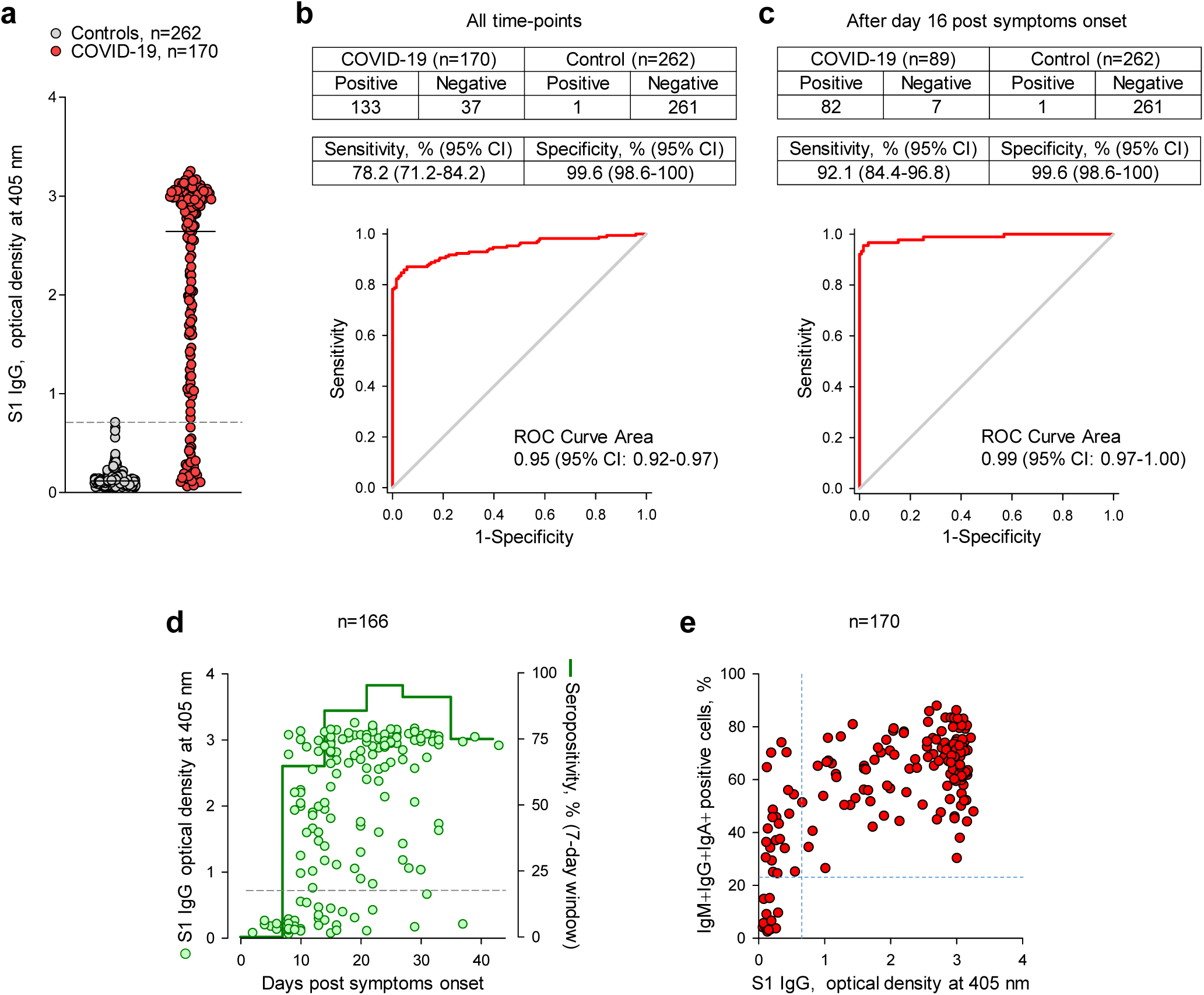
Performance of the S1 ELISA assay for SARS-CoV-2-reactive antibody detection. **a**, Optical densities (ODs) from S1-coated ELISAs on sera from 170 confirmed COVID-19 cases and 262 controls. The dashed line represents the calculate cut-off for positivity. **b-c**, Sensitivity and specificity for the assay calculated for all the sample collected at any time-point post symptoms onset (b) or only for samples collected 16 days after the onset of COVID-19 symptoms (c). The receiver operating characteristics (ROC) curve areas and 95% confidence intervals (CI) are also shown for each group. **d**, Kinetics of seroconversion in COVID-19 patients determined by the S1-coated ELISA. ODs are plotted over time of sample collection since symptoms onset. Seropositivity was calculated for each consecutive week since symptoms onset. Only COVID-19 patients with known date of symptoms onset are included. **e**, Correlation of seropositivity determined by the S1-coated ELISA and the FACS-based assay. Each symbol represents and individual COVID-19 patient sample.

**Extended data Fig. 8.**
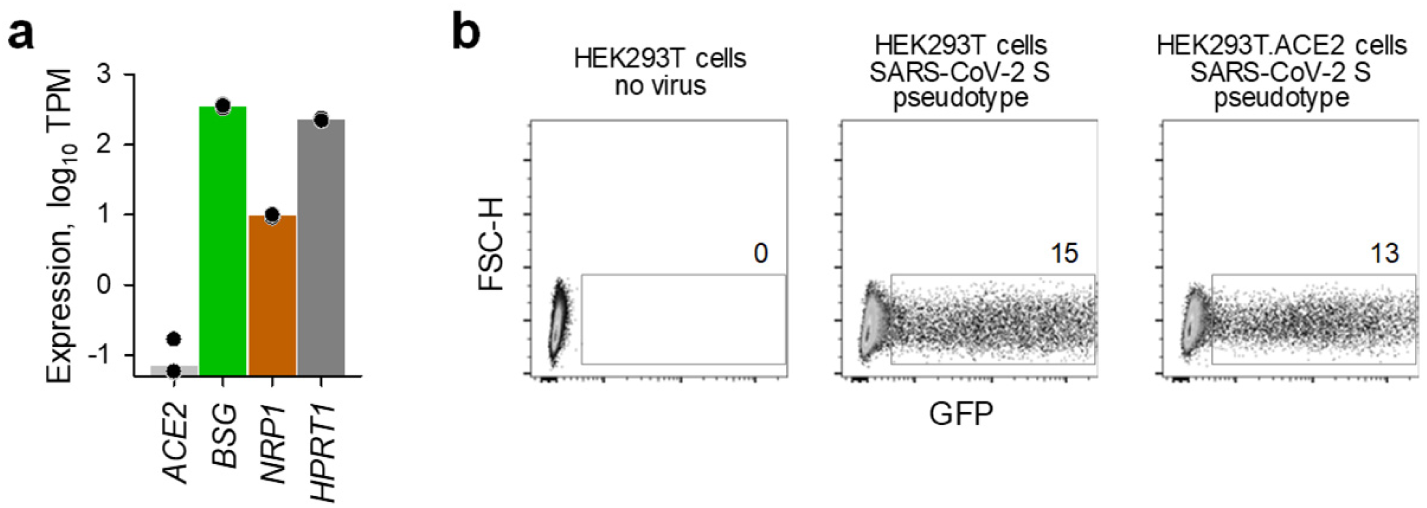
Entry of SARS-CoV-2 S pseudotypes in HEK293T cells. **a**, Expression, plotted as transcripts per million (TPM), of *ACE2, BSG*, (encoding CD147), *NRP1* (encoding neuropilin 1) and *HPRT1* in public RNA-sequencing data (GSE85164) from HEK293T cells. **b**, Transduction efficiency of parental HEK293T cells and HEK293T cells overexpressing ACE2 (HEK293T.ACE2) with GFP-encoding SARS-CoV-2 S pseudotyped lentiviral particles.

**Extended data Fig. 9.**
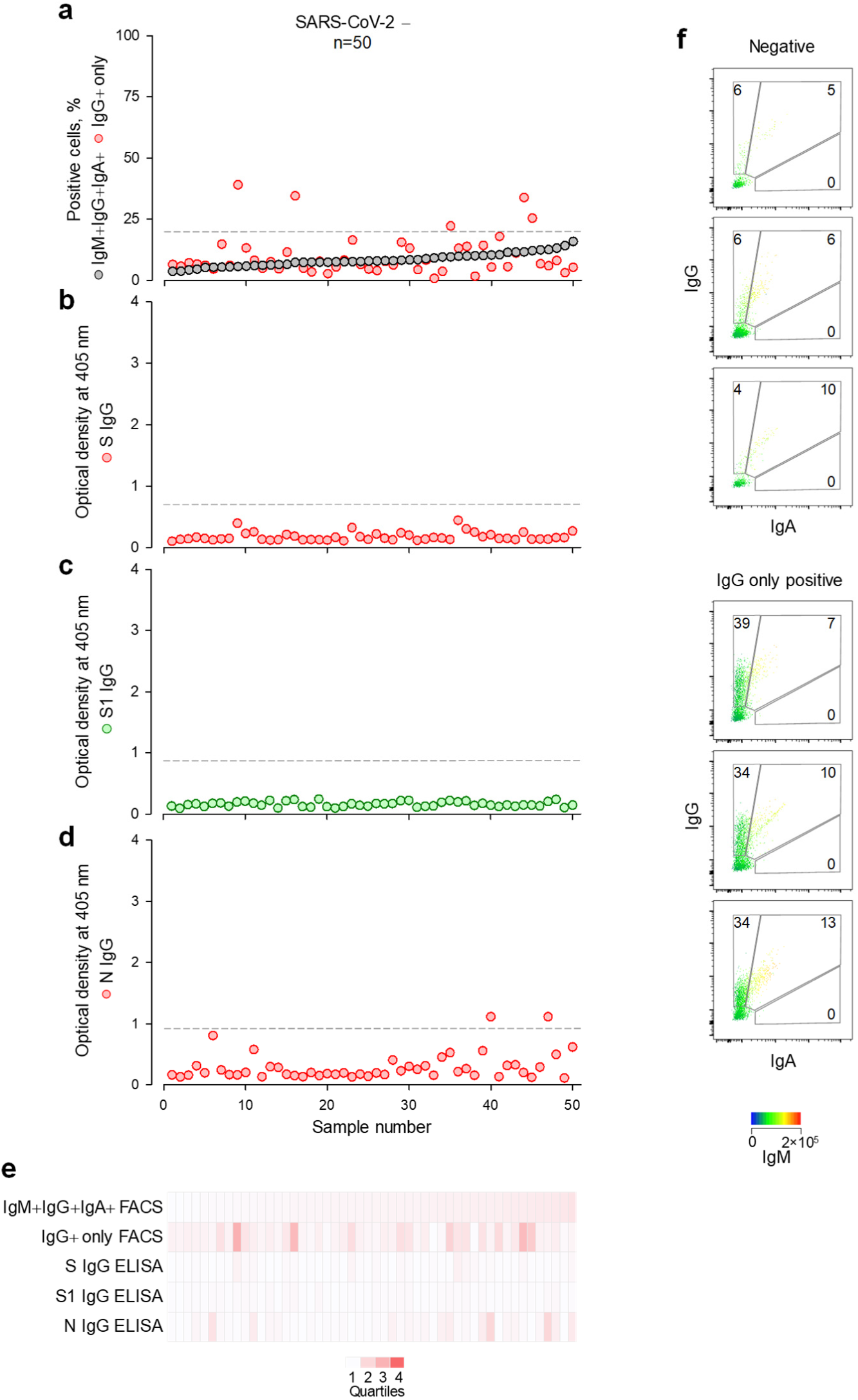
SARS-CoV-2-reactive antibody detection in an additional control cohort. Samples from a total of 50 SARS-CoV-2-uninfected individuals collected in 2018 were tested. All 50 were pregnant healthy women visiting antenatal clinics (Table S1). **a**, Frequency of cells that stained with all three antibody classes (IgM+IgG+IgA+) or only with IgG (IgG+) in each of these samples, ranked by their IgM+IgG+IgA+ frequency. **b-d**, Optical densities from ELISAs coated with S (b), S1 or RBD (c) or N (d) of the same samples. Dashed lines in a-d denote the assay sensitivity cut-offs. **e**, Summary of the results from a-d, represented as a heatmap of the quartile values. **f**, Representative samples that are negative for all Ig classes (Negative) or positive only for IgG (IgG positive).

**Extended data Fig. 10.**
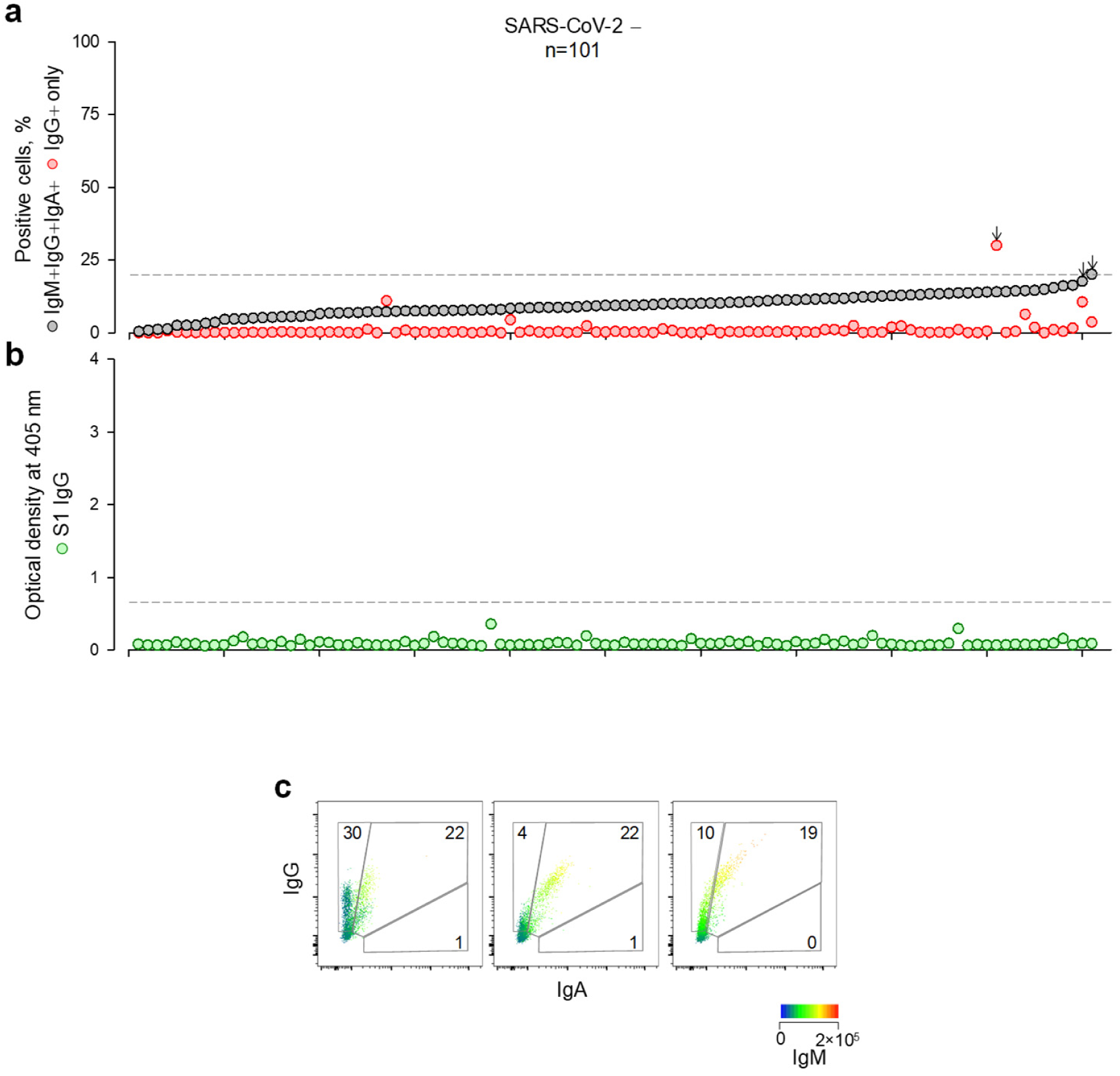
SARS-CoV-2-reactive antibody detection in an additional control cohort. Samples from a total of 101 SARS-CoV-2-uninfected individuals collected in 2019 were tested. These included patients with unrelated viral or bacterial infections (Table S1). **a**, Frequency of cells that stained with all three antibody classes (IgM+IgG+IgA+) or only with IgG (IgG+) in each of these samples, ranked by their IgM+IgG+IgA+ frequency. Arrows indicate the 3 samples with SARS-CoV-2-cross-reactive antibodies. **b**, Optical densities from S1-coated ELISA of the same samples. Dashed lines in (a, b) denote the assay sensitivity cut-offs. **c**, FACS profiles of the 3 samples that were positive for SARS-CoV-2-cross-reactive antibodies.

**Extended data Fig. 11.**
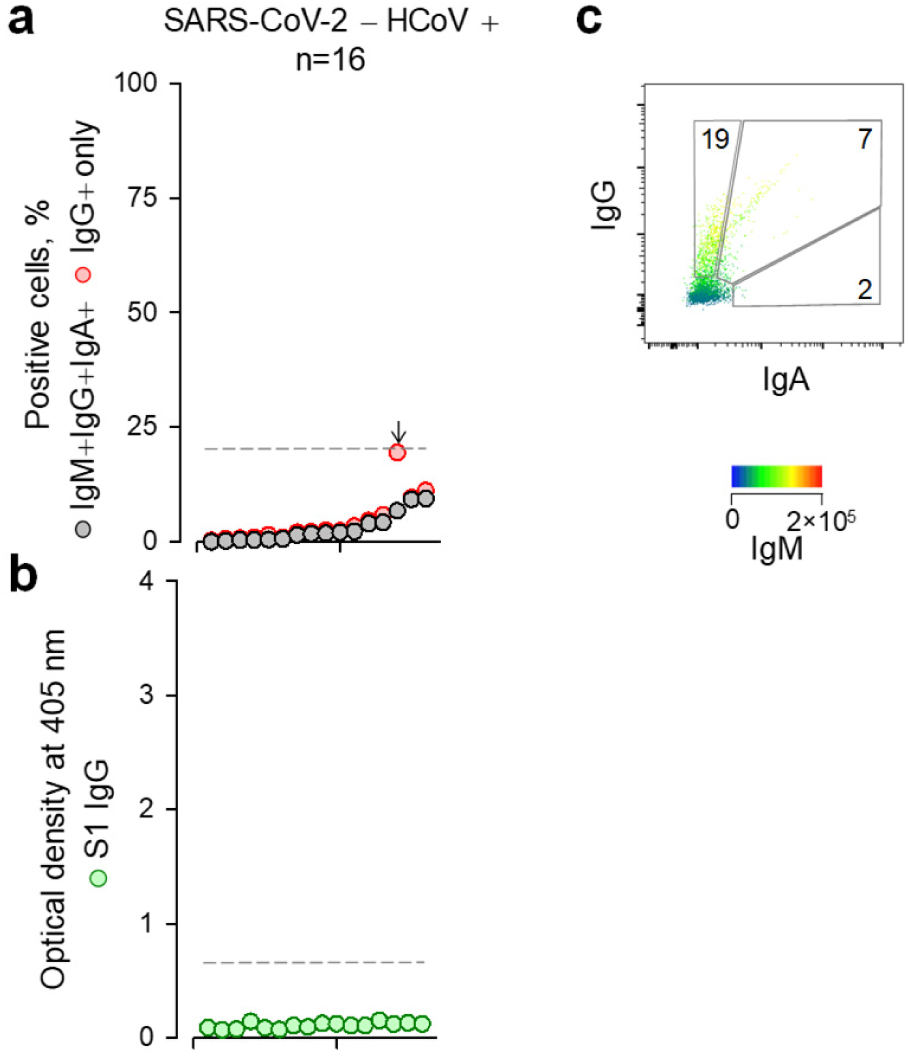
SARS-CoV-2-reactive antibody detection in an additional control cohort. A total of 16 samples from 13 individuals with recent HCoV infection (Table S1) were tested. One haematology patient, persistently infected with NL63 was sampled 4 separate times and all other donors were sampled once. **a**, Frequency of cells that stained with all three antibody classes (IgM+IgG+IgA+) or only with IgG (IgG+) in each of these samples, ranked by their IgM+IgG+IgA+ frequency. The arrow indicate sample with SARS-CoV-2-cross-reactive antibodies. **b**, Optical densities from S1-coated ELISA of the same samples. Dashed lines in (a, b) denote the assay sensitivity cut-offs. **c**, FACS profile of the sample with SARS-CoV-2-cross-reactive antibodies from 66- year-old donor infected with OC43 18 days prior to sampling.

**Extended data Fig. 12.**
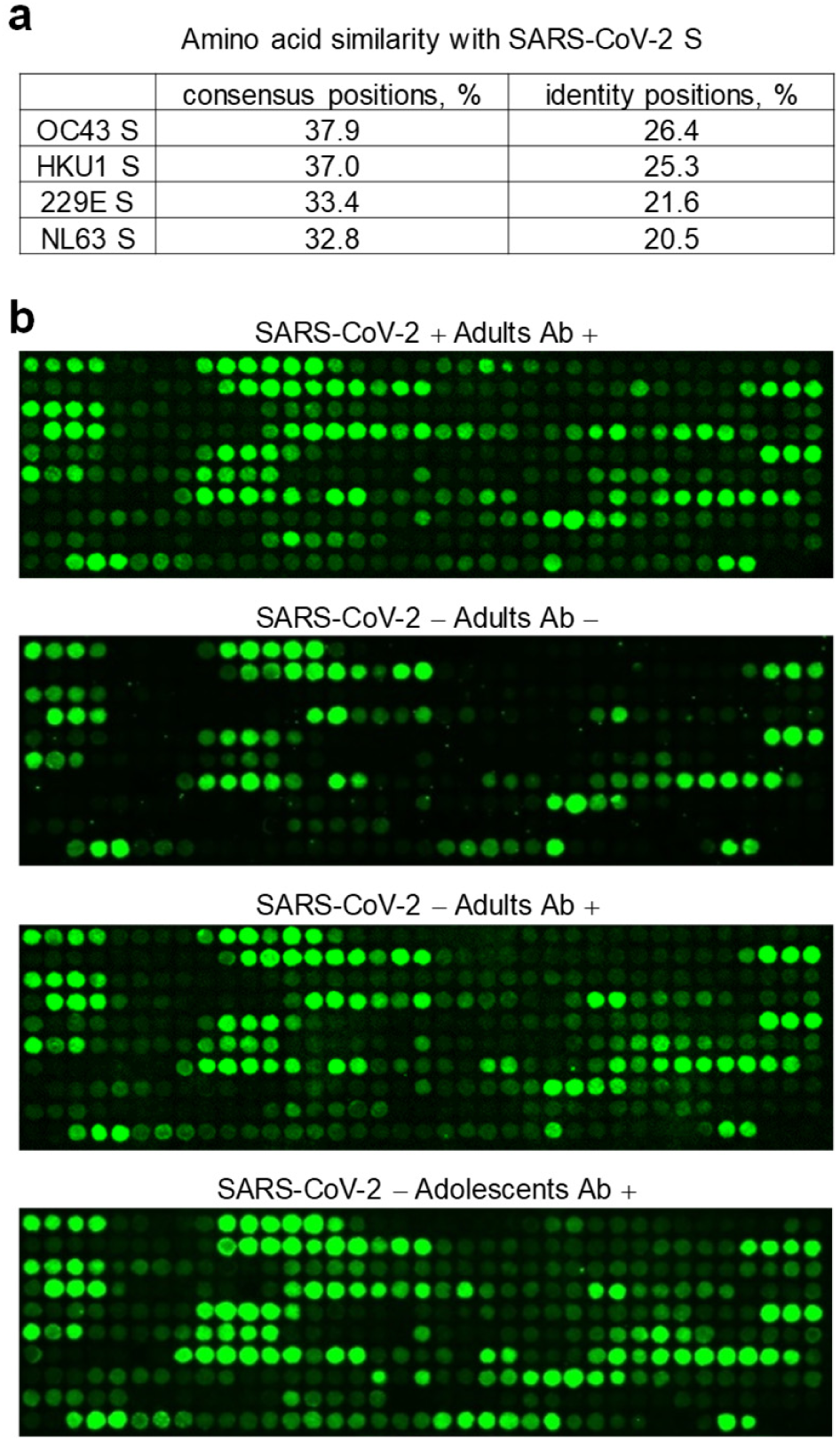
Mapping of cross-reactive epitopes in SARS-CoV-2 S using peptide arrays. **a**, Amino acid sequence similarity and identity between SARS-CoV-2 S and the S proteins of each of the 4 types of HCoV. **b**, Scanned images of peptide arrays spanning the last 743 amino acids of SARS-CoV-2 S detected with primary sera from seroconverted COVID-19 patients (SARS-CoV-2 + Adults Ab +), adult SARS-CoV-2-uninfected donors without cross-reactive antibodies detectable by FACS (SARS-CoV-2 - Adults Ab -), and adult and adolescent SARS-CoV-2- uninfected donors that had cross-reactive antibodies detectable by FACS (SARS-CoV-2 -Adults Ab + and SARS-CoV-2 - Adolescents Ab +, respectively). The signal of the secondary antibody label (IRDye 800CW) is shown in green. The top left position in each array is the first peptide in the sequence (S_531-542_). The 12-mer peptides were arranged from left-to-right and top-to-bottom with an overlap of 10 amino acids, creating 367 spots.

**Extended data Fig. 13.**
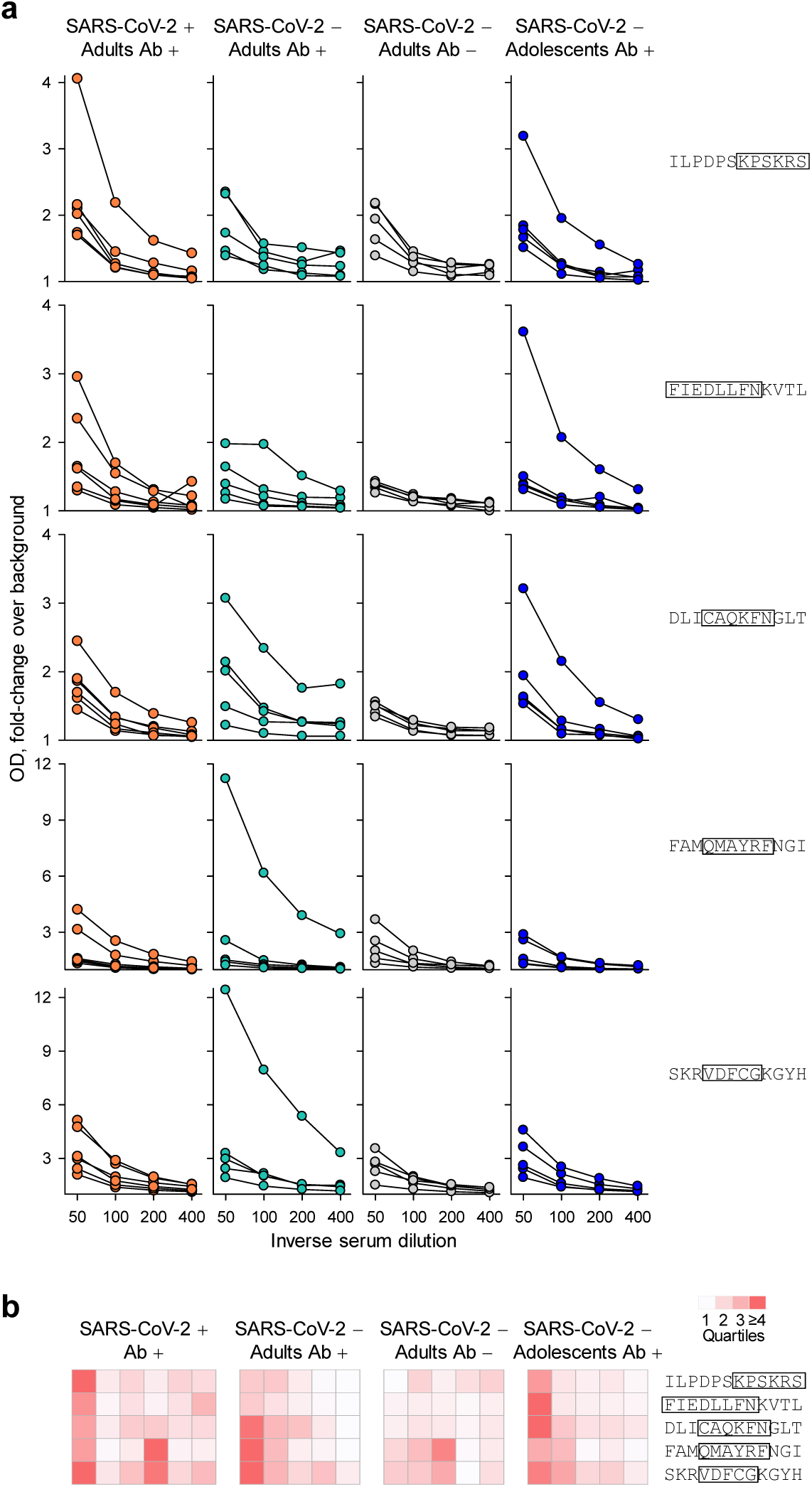
Reactivity against shared peptide epitopes determined by ELISA. Sera from seroconverted COVID-19 patients (SARS-CoV-2 + Adults Ab +, n=6), adult SARS-CoV-2-uninfected donors without FACS-detectable cross-reactive antibodies (SARS-CoV-2 - Adults Ab -, n=5), and adult and adolescent SARS-CoV- 2-uninfected donors that had cross-reactive antibodies detectable by FACS (SARS-CoV-2 -Adults Ab +, n=5 and SARS-CoV-2 - Adolescents Ab +, n=5, respectively), were used in ELISAs coated with the indicated peptides. **a**, Results are shown as fold-change between sample ODs and ODs of negative control wells. Each line represents an individual sample over the indicated serial dilutions. **b**, Summary of reactivity of individual sera from the 4 indicated groups. Each column is an individual sample at 1:50 dilution, represented as a heatmap.

**Extended data Fig. 14.**
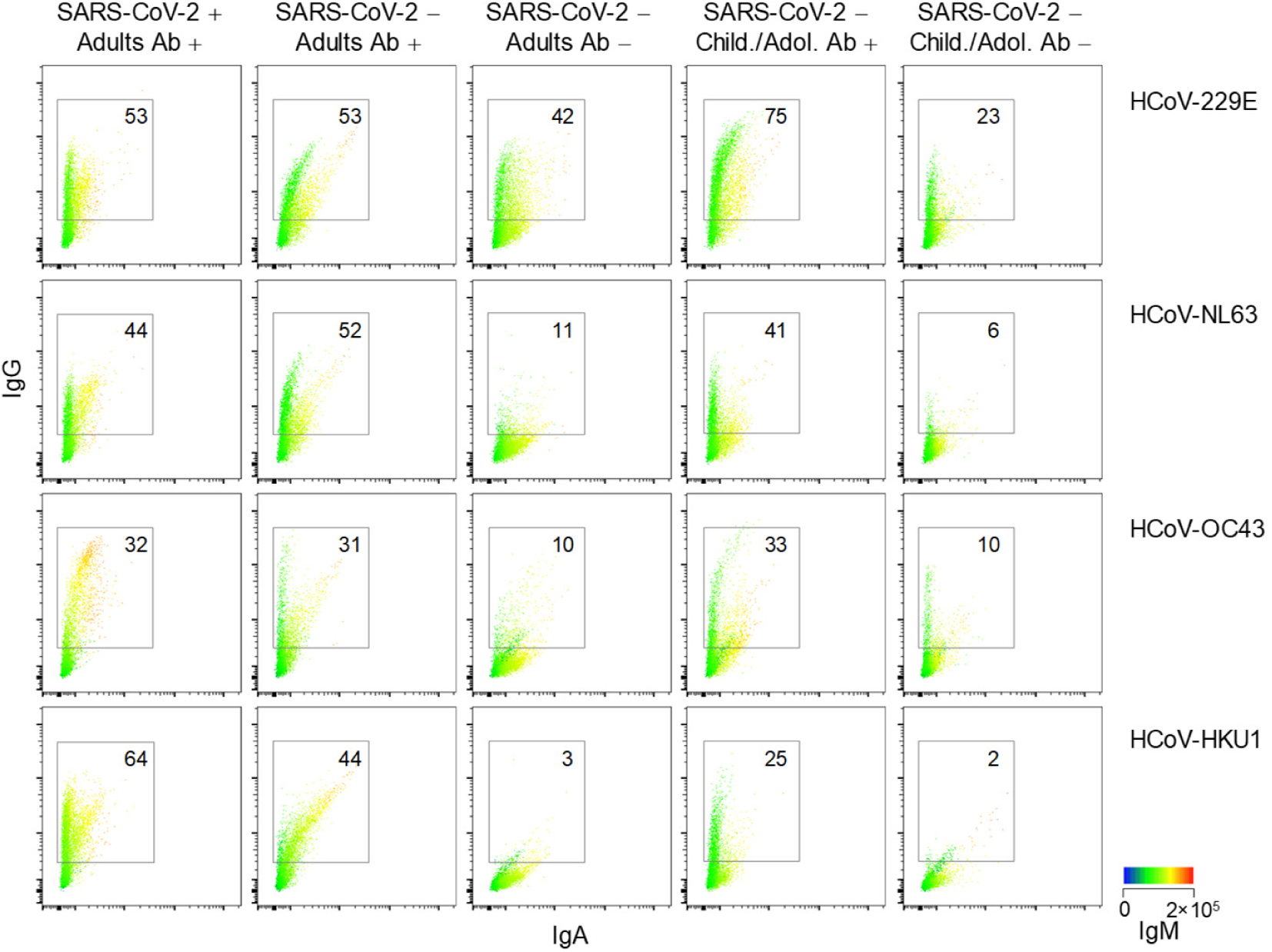
Reactivity against the S glycoproteins of HCoVs determined by FACS. FACS profiles of HEK293T cells transfected to express the S glycoproteins of each of the 4 HCoVs and stained with the indicated sera (at 1:50 dilution). The same representative sample for each group is shown for all HCoVs for consistency. The groups include seroconverted adult COVID-19 patients (SARS-CoV-2 + Adults Ab +); SARS-CoV-2-uninfected adults or children/adolescents with (SARS-CoV-2 - Adults Ab + and SARS-CoV-2 - Child./Adol. Ab +, respectively) or without (SARS-CoV-2 - Adults Ab - and SARS-CoV-2 - Child./Adol. Ab -, respectively) FACS-detectable antibodies cross-reactive with SARS-CoV-2 S. Levels of IgA and IgG are indicates in the x and y axes, respectively, and levels of IgM are indicated by a heatmap. Numbers within the plots denote the percentage of cells stained with IgG antibodies, irrespective of co-staining with IgM or IgA. Those stained with IgM or IgA are not shown here, but are summarised in Fig. 5d.

## Notes

### Competing Interest Statement

The authors have declared no competing interest.

